# Comparative single-cell analyses reveal evolutionary repurposing of a conserved gene program in bat wing development

**DOI:** 10.1101/2024.10.10.617585

**Authors:** Magdalena Schindler, Christian Feregrino, Silvia Aldrovandi, Bai-Wei Lo, Anna A. Monaco, Alessa R. Ringel, Ariadna Morales, Tobias Zehnder, Rose Yinghan Behncke, Juliane Glaser, Alexander Barclay, Guillaume Andrey, Bjørt K. Kragesteen, René Hägerling, Stefan Haas, Martin Vingron, Igor Ulitsky, Marc Marti-Renom, Julio Hechavarria, Nicolas Fasel, Michael Hiller, Darío Lupiáñez, Stefan Mundlos, Francisca M. Real

## Abstract

Bats are the only mammals capable of self-powered flight, an evolutionary innovation based on the transformation of forelimbs into wings. The bat wing is characterized by an extreme elongation of the second to fifth digits and a wing membrane called *chiropatagium* connecting them. Here we investigated the developmental and cellular origin of this structure by comparing bat and mouse limbs using omics tools and single-cell analyses. Despite the substantial morphological differences between the species, we observed an overall conservation of cell populations and gene expression patterns including interdigital apoptosis. Single-cell analyses of micro-dissected embryonic chiropatagium identified a specific fibroblast population, independent of apoptosis-associated interdigital cells, as the origin of this tissue. These distal cells express a conserved gene program including the transcription factors *MEIS2* and *TBX3*, which are commonly known to specify and pattern the early proximal limb. Transgenic ectopic expression of *MEIS2* and *TBX3* in mouse distal limb cells resulted in the activation of genes expressed during wing development and phenotypic changes related to wing morphology, such as the fusion of digits. Our results elucidate fundamental molecular mechanisms of bat wing development and illustrate how drastic morphological changes can be achieved through repurposing of existing developmental programs during evolution.

## Introduction

Evolution has fueled the emergence of a remarkable variety of phenotypes throughout the animal kingdom. In particular, the vertebrate limb displays many fascinating adaptations (*1, 2*) and has long served as a prime example to study the genetic basis of phenotypic evolution (*3, 4*). An extreme example is the evolution of forelimbs (FL) into wings in bats (order *Chiroptera*), the only mammals capable of self-powered flight. Interestingly, the oldest known bat fossil already presents wing-structured FLs, suggesting that flight originated in the most recent common ancestor of all bats (*5*). Bat wings are thus a unique and ancient structure, representing an exceptional model for studying limb diversification. Likewise, examining the development of wings can shed light on the mechanisms underlying morphological transformations in evolution (*6, 7*).

During development, limb buds arise from the lateral plate mesoderm (LPM) under the control of three distinct signaling centers: the zone of polarizing activity, the dorsal and ventral ectoderm, and the apical ectodermal ridge (*8, 9*). These centers confer cellular identity along the anterior-posterior, dorsal-ventral, and proximo-distal axes, respectively. Outgrowth along the proximo-distal axis results in the formation of three distinct elements: most proximally the stylopod, followed by the zeugopod, and distally the autopod, corresponding to humerus/femur, radius-ulna/tibia-fibula and hand/foot, respectively (*10*) (Fig. 1A). The bat forelimb is characterized by elongation of all skeletal elements as well as the presence of membranes which form the wing. Changes are most pronounced in the autopod, with extremely elongated digits II-V and an interdigital wing membrane connecting them, known as chiropatagium. In contrast, in bat hindlimbs (HLs) and most other pentadactyl species including humans and mice, the tissue between the digits recedes during development resulting in separate digits (Fig 1A). Experiments across different species have shown that retinoic acid (RA)-induced apoptosis of interdigital cells plays a central role in digit separation (*11, 12*). Consequently, one hypothesis for the persistence of interdigital tissue in bats is the suppression of this apoptotic process. Several studies have addressed this hypothesis; however, the results have been inconclusive. Both, pro- and anti-apoptotic markers were found to be expressed in the developing chiropatagium (*13, 14*). In addition, several comparative molecular studies have identified genes with altered patterns of expression in developing wings (*15–17*). However, the molecular and evolutionary bases of wing morphology development remain largely unknown, partially due to the limitations of the available methodologies at the time. Recently, single-cell approaches have provided new tools to investigate cell identity and function at unprecedented resolution in any organism, holding great potential to unravel the basis of evolutionary innovation (*18*). Yet, how cell fates are molecularly determined and sustain the emergence of new morphologies remains one of the big unsolved questions in biology.

**Figure 1:**
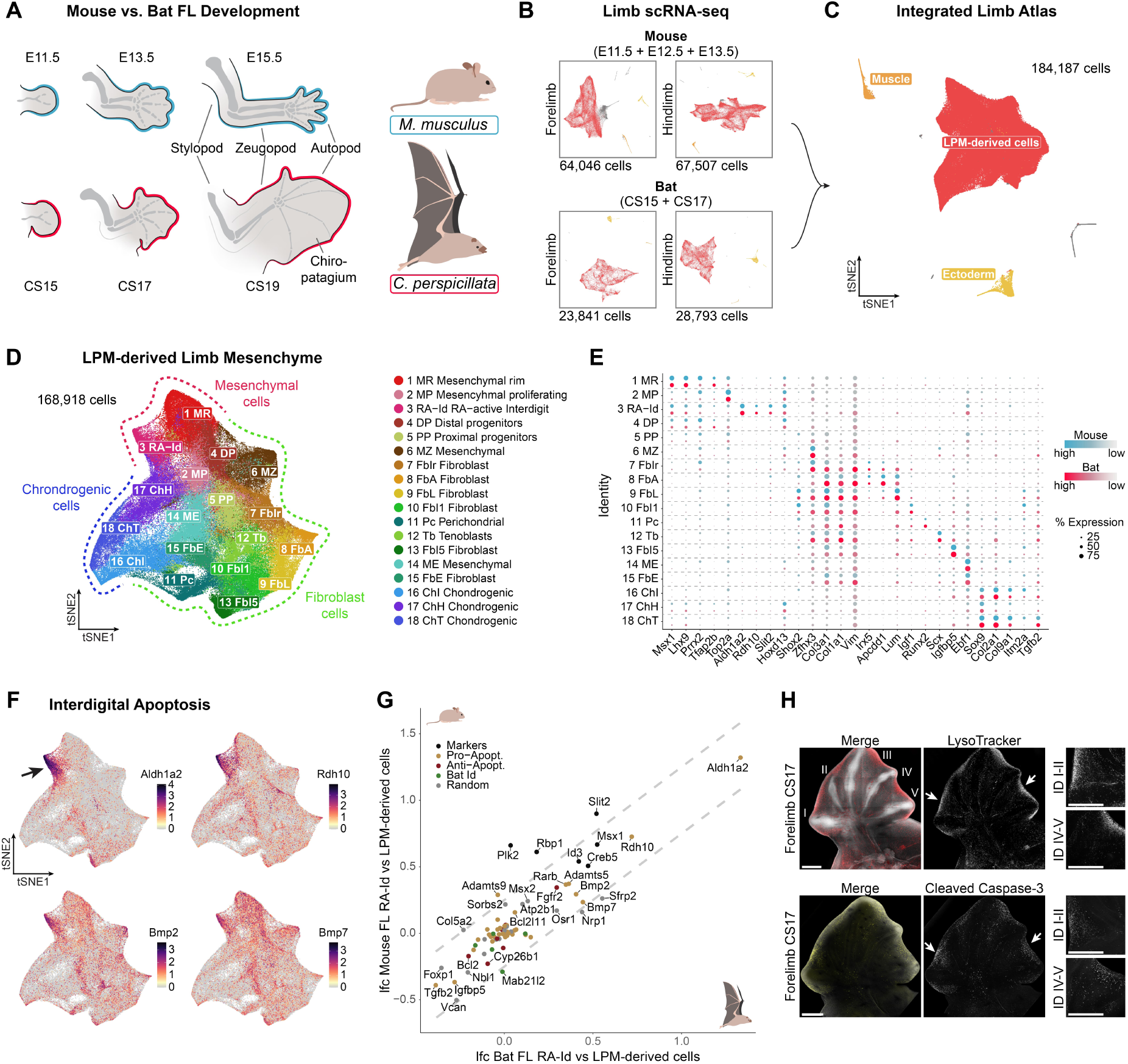
Limb developmental cell states and the process of interdigital apoptotic cell death are conserved in mouse and bat limbs. **A** Schematic representation of key embryonic stages of mouse (blue) and bat (red) limb development. **B** tSNE plots of individual mouse and bat fore- and hindlimb limb single-cell datasets. The main cell-populations of the developing limb are highlighted (red: LPM-derived cells; orange: muscle cells; yellow: ectodermal cells). **C** tSNE plot of integrated inter-species limb atlas. The main cell-populations of the developing limb are highlighted as in B. **D** tSNE plot of sub-clustered LPM-derived cells with cluster annotations. The main developmental lineages are highlighted (red: mesenchymal cells; green: fibroblast cells; blue: chondrogenic cells). **E** Dot-plot showing marker gene expression used for integrated cluster annotation. The color intensity indicates the expression level (blue: mouse; red: bat); the dot size represents the percentage of cells expressing respective marker genes. **F** tSNE plots of the integrated data showing the expression of central components of retinoic acid metabolism and BMP signaling involved in interdigital cell death. The arrow indicates the retinoic acid active interdigital cell population (3 RA-Id). **G** Correlation of pro- (yellow) and anti-apoptotic (red) genes in the 3 RA-Id cell population of mouse and bat. Marker genes of this cell population are highlighted in blue; genes previously reported to be expressed in bat interdigital regions are highlighted in green. Shown is the log fold change (lfc) of differential gene expression analysis between the RA-Id cluster 3 versus the rest of the LPM-derived mesenchyme per species in the FL. A set of random genes was included as control. **H** LysoTracker staining (upper panel) and immunostaining against Cleaved Caspase-3 protein (lower panel) of bat FL at stage CS17 with magnification of interdigital regions (indicated by arrows) between digits I and II (which later lack interdigital membrane) and IV and V (later connected by chiropatagium) shown on right. Merged images show DAPI and LysoTracker or Cleaved Caspase-3 signal. Scale bars represent 500 µm.

## Results

### Cellular composition and interdigital cell death are conserved between bat and mouse limbs

We collected FLs and HLs for single-cell transcriptomics (scRNA-seq) from mice and bats (*C. perspicillata*) covering critical developmental stages of digit separation and wing formation. Samples included an early, morphologically undifferentiated stage (embryonic day (E)11.5 in mice and equivalent to CS15 stage in bats (*19*)) and a later stage in which the digits form and separate (E13.5 in mice and CS17 in bats); we also included an intermediate time point (E12.5) from mice (Fig. 1B). Using the Seurat *SCTransform* integration tool, we generated an inter-species single-cell transcriptomics limb atlas (Fig. 1C). Cells from both species contributed similarly to all cell clusters (Supplementary Fig. 1). We identified all major cell populations known to be present in developing limbs, including muscle, ectoderm-derived and LPM-derived cells (*20–22*) (Fig. 1C, Supplementary Fig. 1). Overall, both the composition and identity of limb cells are largely conserved between the species despite significant morphological differences.

As the LPM contributes to the formation of interdigital mesenchyme, cartilage, tendons, and other connective tissues within the limb, we specifically focused on this lineage. The LPM-derived cells were further subdivided into 18 clusters and annotated by performing differential gene expression analysis. Based on the calculated markers and previous studies (*23, 24*), we identified three main cell lineages: chondrogenic, fibroblast and mesenchymal (Fig. 1D). The expression of top marker genes per cell cluster was also conserved across species (Fig. 1E).

Using this inter-species single-cell atlas, we first sought to address the prevailing hypothesis that chiropatagium development is driven by inhibition or reduction of apoptotic cell death in the interdigital tissue (*13*). We identified a cluster of interdigital cells characterized by high expression levels of *Aldh1a2* and *Rdh10*, components of RA signaling. RA is regarded as a pivotal regulator of interdigital apoptosis and its expression pattern has been extensively employed to discern the interdigital tissue (*25*). Cells from this cluster (3 RA-Id) also expressed main pro-apoptotic factors, including *Bmp2* and *Bmp7*, highlighting it as a central population of apoptotic signaling in both species (Fig. 1F) (*26*). Within this cluster, we then analyzed the expression of a larger number of genes associated with different cell death processes such as *Bcl2*-, *Bmp*- and *Fgf*-associated signaling and senescence (*27*). Our data revealed no significant relative transcriptional differences in pro- or anti-apoptotic factors for the RA-Id cluster between species (Fig. 1G and Supplementary Fig. 2). Interestingly, genes known to be distinctively expressed in the interdigital tissue of bat wings, including the anti-apoptotic *Grem1* (*13*), also did not show a difference in relative expression, suggesting expression in a different cluster.

To further confirm the presence of apoptosis in bat FLs, we stained bat limbs with LysoTracker, a marker of lysosomal activity which correlates with cell death (*28*). We found pronounced staining in bat FLs at the edges of all interdigital zones, with no discernable differences to interdigits I-II in FLs or any HL interdigit, which all regress (Fig. 1H and Supplementary Fig. 2). In addition, we confirmed that cell death in bat wings occurs via an apoptotic process activated by the caspase cascade, as indicated by the positive staining for cleaved Caspase-3 protein (Fig. 1H and Supplementary Fig. 2). In summary, our analysis revealed that the cell composition between mouse and bat limbs is highly conserved. Furthermore, cell death occurs in interdigital cells in both species.

### The developmental origin of the bat chiropatagium

Since cell death occurs similarly in both bat and mouse interdigital clusters, it cannot account for the persistence of interdigital tissue. To identify the cells that persist and form the chiropatagium, we independently clustered the mouse and bat datasets and compared them with the integrated results. The clusters showed a good correspondence, with a high correlation of gene expression between species (Fig. 2A, B and Supplementary Fig. 3), suggesting that the chiropatagium is not associated to the emergence of a novel cell cluster in the bat wing.

**Figure 2:**
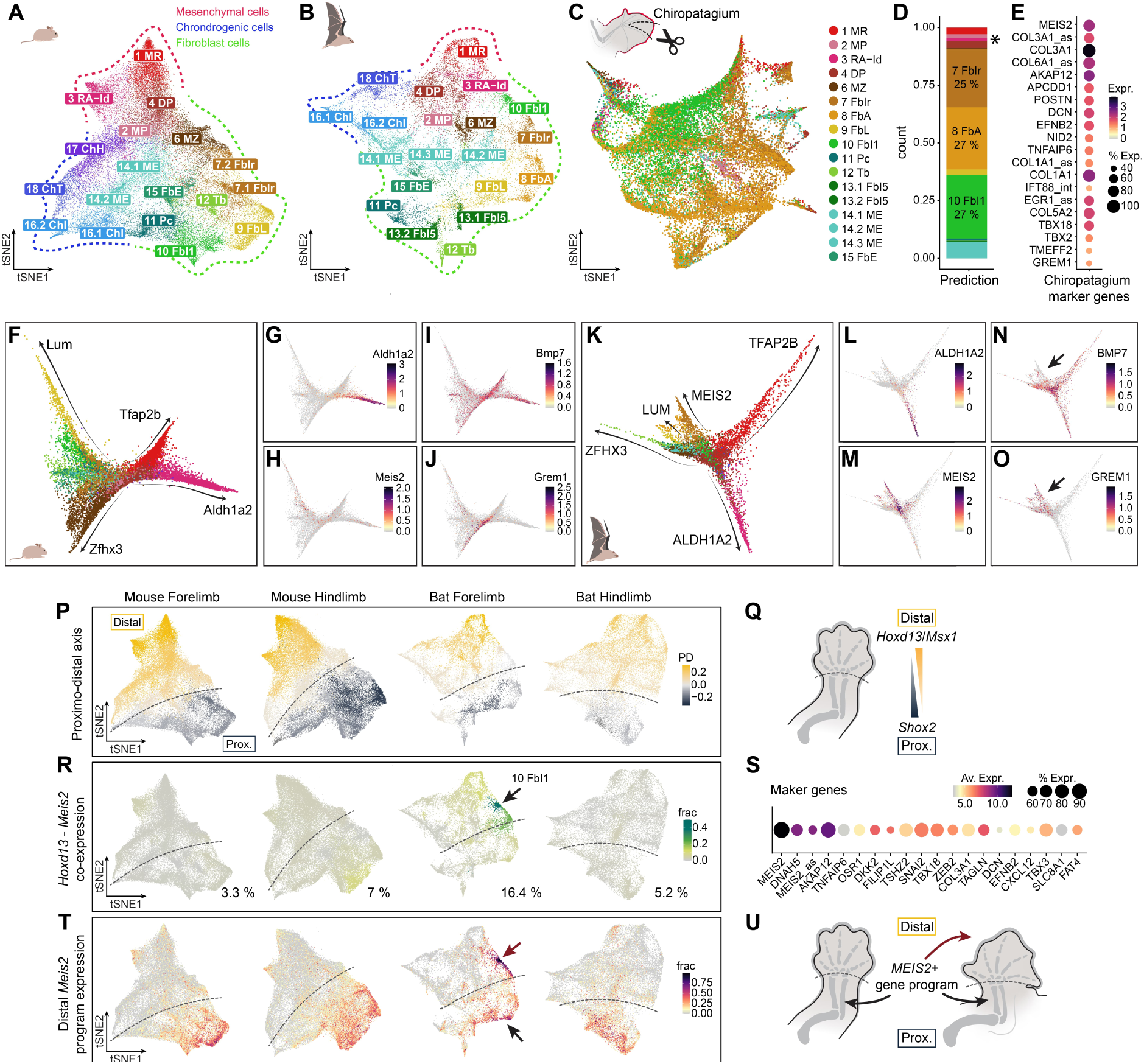
The chiropatagium is composed of fibroblasts highly expressing *MEIS2* following a unique developmental trajectory in the bat forelimb autopod. **A** and **B** tSNE plots of individual clustering of mouse (A) and bat (B) FL data. The main developmental lineages are highlighted (red: mesenchymal cells; green: fibroblast cells; blue: chondrogenic cells). The cluster labels and colors are derived from Figure 1D. **C** tSNE plot of micro-dissected Chiropatagium cells from CS18 bat FLs. Cells are colored and labeled by their transcriptional correspondence to bat FL LPM-derived mesenchyme (B). **D** Quantification of cluster correspondence between chiropatagium cells and LPM-derived mesenchymal cells. RA-Id cells are indicated by an asterisk. **E** Marker genes of chiropatagium cells based on differential gene expression of chiropatagium cells versus bat FL LPM-derived mesenchymal cells. **F** Differentiation trajectories of *Hoxd13*- positive, non-chondrogenic cells of the mouse FL, derived from RNA velocity and pseudotime data indicated by arrows. Trajectories were annotated based on increasing expression of marker genes. **G-J** Expression of *Aldh1a2*, *Meis2*, *Bmp7* and *Grem1* in mouse FL trajectories. **K** Differentiation trajectories of *HOXD13*-positive, non-chondrogenic cells of the bat FL, derived from RNA velocity and pseudotime data indicated by arrows. Trajectories were annotated based on increasing expression of marker genes. **L-O** Expression of *ALDH1A2*, *MEIS2*, *BMP7* and *GREM1* in bat forelimb trajectories. The arrow in N indicates a unique *MEIS2*-positive trajectory identified in the bat FL. **P** Assignment of a proximal (dark blue) or distal (yellow) identity to each cell of mouse and bat fore- and hindlimbs based on *Hoxd13*+*Msx1* and *Shox2* expression per cell. Shown are the mean differences of the expression scores per cluster. **Q** Schematic representation of distal and proximal markers used in P. **R** Co-expression of distal autopodial marker *Hoxd13* and chiropatagium marker *Meis2* in mouse and bat fore- and hindlimbs. Shown is fraction of cells co-expressing *Hoxd13* and *Meis2* per cluster. Cluster 10 FbI1 is highlighted with an arrow. **S** Marker genes of bat cluster 10 Fbl1 based on differential gene expression between cluster 10 and the rest of the LPM-derived bat FL cells. Ordered by adjusted p-value. **T** Bat cluster 10 Fbl1 gene set expression in mouse and bat fore- and hindlimbs. Shown is the co-expression score of the gene set shown in S. The distal and proximal cells highly expressing this program are indicated with an arrow. **U** Schematic representation of the spatial expression of the identified *MEIS2*-positive cell program common in mouse and bat proximal fibroblast cells, while its distal expression is wing-specific.

To trace down the molecular and cellular nature of the chiropatagium, we performed scRNA-seq from micro-dissected bat interdigital tissues at a later stage (CS18, equivalent to E14.5 in mice) (Fig. 2C). We annotated the chiropatagium LPM-derived populations by label transfer using the bat FL LPM data as reference (*29, 30*). This revealed that the chiropatagium is primarily composed of three different populations of fibroblast cells, with transcriptional correspondence to clusters 7 FbIr, 8 FbA, and 10 FbI1 (Fig. 2D). Differential expression analyses against the whole FL LPM dataset showed that the chiropatagium features high expression of *MEIS2*, *COL3A1*, *AKAP12*, and *GREM1* among others (Fig. 2E). Notably, the RA-Id cluster 3 was minimally represented in the chiropatagium (∼ 1%, Fig. 2D), which is consistent with the results of the apoptosis staining (Fig. 1H). Thus, the 3 RA-Id cluster can be ruled out as the cellular source of the chiropatagium.

To further elucidate the origin of chiropatagium cells, we inferred developmental trajectories in mouse and bat distal LPM clusters, focusing on non-skeletal cells expressing *Hoxd13*, a *bona fide* marker of the autopodial lineage (*31–33*). Using the RNA velocity tool *scVelo*, as well as the pseudotime tool *slingshot*, we identified independent trajectories that share the same origin and are defined by differential increased gene expression (Fig. 2F-O and Supplementary Fig. 4). For example, the 3 RA-Id cluster forms a trajectory with increasing *Aldh1a2* expression (Fig. 2G, L). Moreover, in bat FL we identified an independent trajectory of fibroblasts marked by the expression of the transcription factor (TF) *MEIS2*. This trajectory was neither detected in mice nor in bat HLs, suggesting a unique developmental specification for chiropatagium cells (Fig 2H<u>, M</u> and Supplementary Fig. 4). Overall, these analyses further show that the chiropatagium develops independently from the 3 RA-Id interdigital population. In contrast, this tissue is primarily composed of fibroblast cells expressing *MEIS2*.

MEIS2 is a TF that defines proximal identity at early limb stages (*34, 35*). To explore its distal role during bat autopod morphogenesis, we first defined the proximo-distal identity for each cell and cluster across all non-integrated datasets. Specifically, we calculated the gene expression ratio between distal (autopod) and proximal markers (*Hoxd13* + *Msx1* vs *Shox2*). Most clusters could be clearly identified as either proximal or distal (Fig. 2P, Q). *Meis2* was among the marker genes characterizing the proximal non-skeletal cells in all our samples (Supplementary Fig. 3 and 4). We then quantified the fraction of *Meis2*-positive cells in the distal region by calculating the co-expression of *Hoxd13* and *Meis2*. This analysis revealed that the highest number of cells expressing both factors and highest co-expression levels are found in the bat FL (Fig. 2R, green color, 16.4%). This co-expression pattern specifically highlighted fibroblast cluster 10 (arrow in Fig. 2R), followed by cluster 7, each one constituting ∼1/3 of chiropatagium cells at later stages (Fig. 2D). We therefore focused on cluster 10 and, by comparing it against the remaining LPM cells, identified 20 marker genes including the TFs *OSR1*, *TBX18* and *TBX3* (Fig. 2S). Given the unusual nature of this cluster, with many cells highly co-expressing distal and proximal markers, we explored the expression of these 20 genes across all samples. Intriguingly, this gene set was found co-expressed at high levels in the proximal fibroblasts (mostly cluster 8 and 9) of FLs and HLs of both species, while its distal co-expression was unique to the bat FL (Fig. 2T). Similar results were found for the marker genes of cluster 7 (Supplementary Fig. 5). Thus, the chiropatagium consists of fibroblasts that do not derive from the RA-Id cells. Rather, chiropatagium cells display their own differentiation trajectory characterized by a specific set of genes that include *MEIS2*, a TF expressed prominently in the proximal limb (Fig. 2U).

### Repurposing of a proximal gene program in the distal bat forelimb

Our analyses identified a fibroblast cluster that is unique to the distal bat FL, yet expresses a gene set that is also present in proximal fibroblast cells of mouse and bat limbs. To determine the degree of transcriptional similarity among these clusters, we performed differential gene expression analyses in bat FLs comparing the proximal (8) and distal (10) fibroblasts against the rest of the LPM cells. We found 223 overexpressed genes, 65% of them (144) displaying high relative expression in both proximal and distal clusters (Fig. 3A). Nevertheless, a subset of genes was specific to distal or proximal clusters (25 and 64, respectively) (Fig. 3B). Interestingly, 34 of the shared genes were also highly expressed in mouse proximal fibroblasts, suggesting an evolutionary conserved function for this gene set in limb fibroblasts (triangular points in Fig. 3A, C). Thus, the distal *MEIS2*-positive cluster 10 is characterized by a gene program that shows significant transcriptional overlap with a proximal cluster. Gene ontology (GO) enrichment analysis for the shared genes revealed distinct functions, including mesenchymal proliferation, extracellular matrix (ECM) organization, and ameboidal-type cell migration (Fig. 3D). These processes are not only indicative of fibroblast identity (*36*), but also represent essential components of interdigital remodeling (*37*) and may be highly relevant in the context of wing development. To better understand the relationship and hierarchies of the genes in the program, we performed a gene regulatory network analysis using *SCENIC* for each cell cluster. This analysis placed MEIS2 in the regulon with the highest regulatory score within the bat cluster 10 (RSS > 0.23; Supplementary Fig. 7). Furthermore, MEIS2 also appeared as a direct regulator for numerous genes, including several that belong to the shared proximo-distal program (Fig. 3E and Supplementary Fig. 7C).

**Figure 3:**
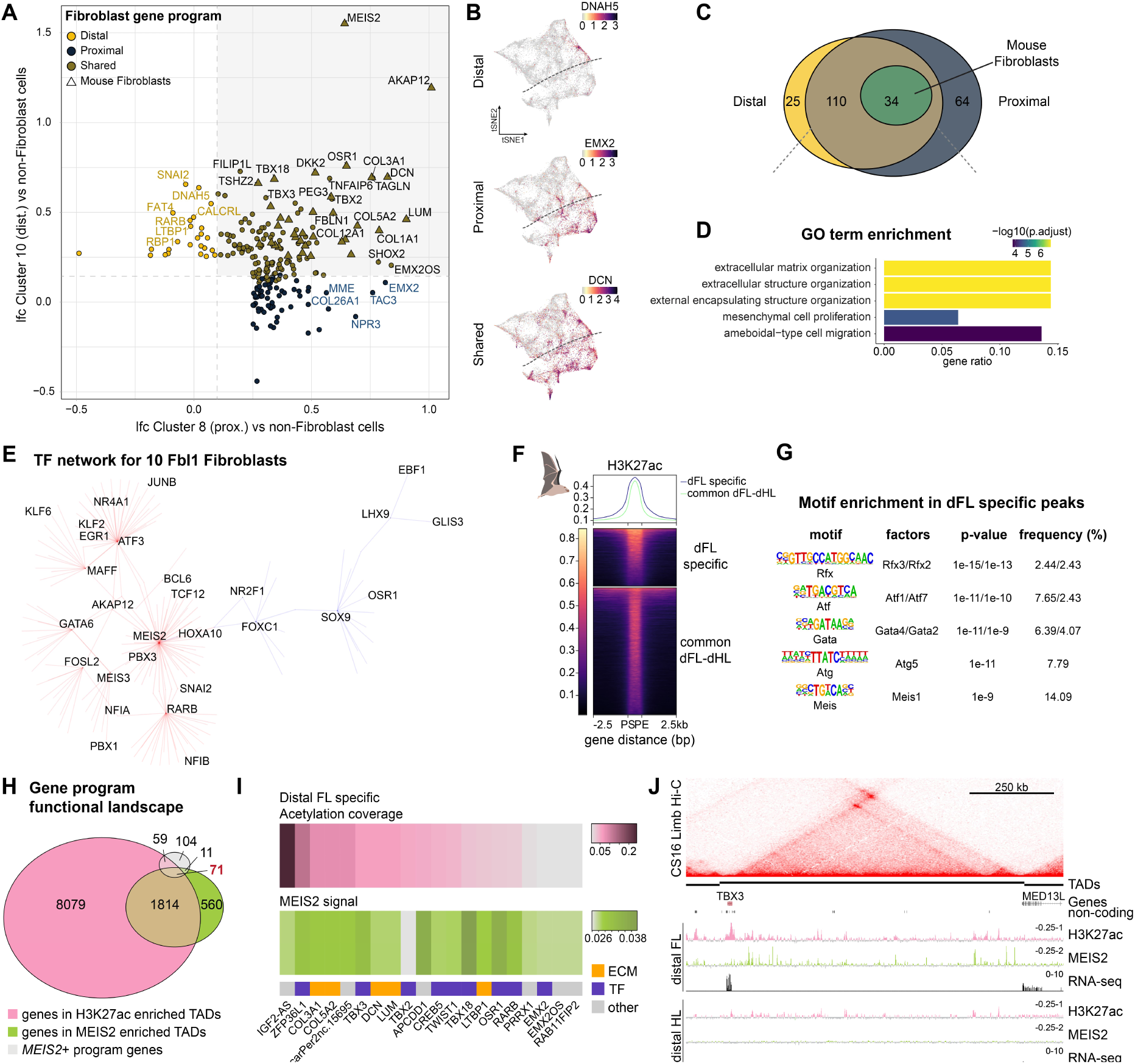
Proximo-distal dissection of mouse and bat limbs reveals repurposing of a proximal gene program in the distal bat forelimb. **A** Correlation between the differential expression of genes from distal (cluster 10) and proximal (cluster 8) *MEIS2*- positive clusters in the bat FL identified in Figure 2. Shown is the lfc of differential gene expression analysis of the respective cluster versus non-fibroblast LPM-derived cells. Genes shared with mouse fibroblasts are depicted as triangles. **B** Representative tSNE plots of genes expressed in the distal, proximal or both clusters of the bat forelimb. **C** Venn diagram showing the overlap (brown) between the genes enriched in the proximal (dark blue) and distal (yellow) cell subset as well as the fraction of genes shared with mouse fibroblasts (green). **D** GO term enrichment analysis of the 144 shared genes. Shown are the top 5 enriched GO terms. E *SCENIC* TFs network analysis for genes enriched in cluster 10. Red lines represent positive regulatory connections, while blue lines indicate negative regulatory connections. **F** Tornado plot showing H3K27ac peaks specific to the distal forelimb as well as common peaks of distal fore- and hindlimb. Shown are the regions from peak start (PS) to peak end (PE). **G** Motif enrichment in distal forelimb specific H3K27ac peaks. Shown are the top 5 binding motifs per TF family. **H** Venn diagram showing the overlap between genes in H3K27ac-enriched and MEIS2-binding enriched TADs, as well as genes from the fibroblast gene program from A. **I** Heatmaps showing the portion of each TAD covered by acetylation peaks, and the mean signal per TAD of MEIS binding. Shown are the top 20 genes by MEIS binding signal. **J** Bat *TBX3* locus with Hi-C from CS16 FLs on top, TAD calling below. The Input subtracted H3K27ac ChIP-seq track is depicted in pink and input subtracted MEIS2 ChIP-seq track is shown in green. RNA-seq tracks are shown in black. ChIP-seq and RNA-seq were performed on distally dissected fore- and hindlimbs at CS18/19.

To further elucidate how this gene program is regulated, we generated bulk transcriptomic and epigenomic datasets from distal limbs by physically dissecting them at the level of the wrist (mouse E15.5 and bat CS19 stages) (Supplementary Fig. 8). Differential expression analyses between distal FL and distal HL showed only small differences for mice, while bat distal FLs showed a higher number of differentially expressed genes (DEGs) compared to HLs. Among the most upregulated genes we found the TFs *MEIS2*, *HOXD9*, *HOXD10*, *HOXA2* and *TBX3*, genes known to be early proximal markers and patterning factors (*38, 39*) (Supplementary Fig. 8). Differential enrichment analyses for active epigenomic regions (marked by H3K27ac) revealed a significant number of regions specific to the distal bat FL, enriched in TF binding sites for RFX, ATF, GATA, ATG and, notably, MEIS (Fig. 3F, G and Supplementary Fig. 8). Since several analyses suggested that MEIS2 plays a critical role in chiropatagium development, we profiled its chromatin binding in distal bat limbs using a dual antibody ChIP-seq assay (*34*). As TFs usually bind to several enhancer regions across large genomic distances (*40*), we summed up all MEIS2-bound regions per regulatory domain, defined by genome-wide chromatin interaction maps (Hi-C) from developing bat limbs. We identified a subset of regulatory domains distinctly enriched with MEIS2 binding signal (Supplementary Fig. 8). By intersecting H3K27ac- and MEIS2-binding enriched domains with genes from the distal/proximal fibroblast gene program, we narrowed down the list of candidate genes potentially regulated by MEIS2 to 71 (Fig. 3H). The top 20 genes displaying the highest overall MEIS2 binding signal in their regulatory domains included genes from the fibroblast gene program, like ECM components and TFs such as *TBX3* and *TBX18* (Fig. 3I). The striking pattern of chromatin activity profiles (H3K27ac and MEIS2 binding) being constrained within regulatory domains is exemplified for the *TBX3* domain (Fig. 3J, and TBX2 in Supplementary Fig. 8). In addition, we compared MEIS2 binding in the distal bat limb with ChIP-seq data from early (E10.5) mouse embryonic limbs (*34*), where MEIS1/2 is known to have a crucial role in limb patterning. The limited overlap in bound gene promoters (21 regions) suggests that MEIS2 has a distinct regulatory role and differential genome accessibility at both stages (Supplementary Fig. 8). In summary, we identify MEIS2 as a critical TF regulating chiropatagium development, through the pervasive binding at the chromatin landscape of its associated gene program.

### Distalization of *MEIS2* and *TBX3* expression induces developmental processes related to wing formation

Our previous analyses positioned *MEIS2* and *TBX3* as key regulators of the gene program associated with chiropatagium development. To investigate their effects on limb developmental cell states, we induced the distal limb expression of these two TFs in transgenic mice. The sequences of these TFs in both species result in highly similar proteins (Supplementary Fig. 9). Constructs were generated in which the bat coding sequences of *MEIS2* and *TBX3* were expressed under the control of a previously characterized *Bmp2* enhancer (*41*). This enhancer has specific activity in the distal non-skeletal mesenchymal and interdigital part of the limb bud (Fig. 4A). This precise spatio-temporal activity allowed the expression of these factors without inducing detrimental effects in other tissues. *In situ* hybridization as well as bulk transcriptomic analysis of mutant limbs at E12.5 validated specific expression of the transgenes in distal mouse limbs (Fig. 4B, Supplementary Fig. 9).

**Figure 4:**
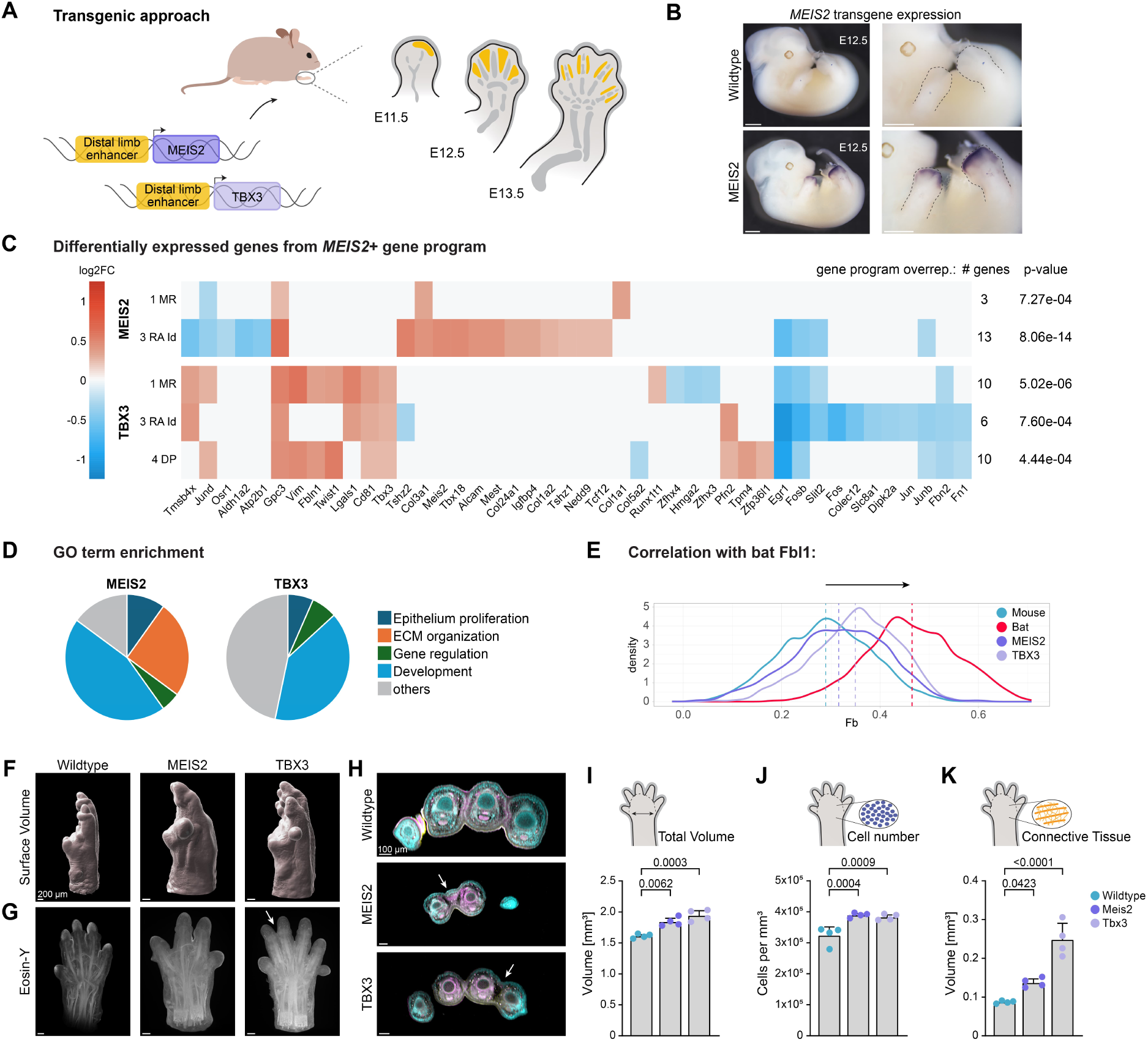
Distal expression of *MEIS2* and *TBX3* in transgenic mouse limbs induces cellular and morphological features related to the chiropatagium tissue. **A** Schematics of transgenics experiments. Bat *MEIS2* and *TBX3* coding sequences were expressed in distal mouse limbs using a characterized *Bmp2* enhancer (*41*) with specific activity in the distal and interdigital mesenchyme (schematically shown in E11.5-E13.5 mouse limbs in yellow). **B** WISH from wildtype and mutant embryos at E12.5 showing distal activity of transgene constructs. n=2. The scale bars represent 1 cm. **C** Expression heatmaps showing differentially expressed genes from the gene program in affected limb clusters of mouse mutant limbs at E12.5 (1 MR and 3 RA Id in *MEIS2* mutant; 1 MR, 3 RA Id and 4 DP in *TBX3* mutant). The number of genes from the fibroblast gene program and p-value from fibroblast gene program over-representation are shown on the right. **D** Pie charts showing enriched GO terms in affected cell clusters. GO terms are grouped by biological functions. **E** Correlation of the affected mutant cells to the bat cluster 10 FbI1, based on the expression of the genes from the gene program. Depicted is the density of the correlation of all cells in the affected clusters, and corresponding clusters in mouse wt and bat forelimb. **F** and **G** 3D imaging of mouse wildtype and mutant limbs at E15.5. Shown is a surface representation (F) and a cross-section showing Eosin-Y staining (G) with an arrow highlighting syndactyly of digit II and III in the *TBX3* mutant. The scale bars represent 200 µm. **H** Cross-sections of the limbs with arrows indicating tissue growth between the digits **I-K** Quantification of autopod surface volume, cell number and connective tissue volume in wildtype and mutant limbs. n=4. Error bars show the standard deviation.

We evaluated the impact on gene expression at cellular resolution by performing scRNA-seq on the mutant limbs at E12.5. Focusing on distal clusters, we isolated *Hoxd13*-positive cells and integrated them with corresponding data from our reference mouse atlas. Using this approach, we performed a differential expression analysis on the clusters where *MEIS2* or *TBX3* were differentially expressed in the mutant samples (Fig. 4C). We found that the genes of the chiropatagium gene program were significantly overrepresented within the DEGs (Fig. 4C). GO enrichment analysis showed that the upregulated genes are involved in ECM production and proliferation processes, functions also characteristic of the identified gene program (Fig. 4D). Moreover, we compared the transcriptomic correlation of mouse wildtype and mutant cells to the mean gene expression of fibroblast cluster 10 from bat FLs. This revealed that mouse mutant cells exhibit higher similarity to these bat cells (Fig. 4E). These results highlight the ability of these two TFs, *MEIS2* and *TBX3*, to partially induce the specific gene program of the chiropatagium.

To evaluate the phenotypic consequences of ectopic distal *MEIS2* and *TBX3* expression, we performed 3D imaging of mutant and wildtype control limbs at a later developmental stage (E15.5). We marked the nuclei with DRAQ5 and used Eosin-Y as a proxy to quantify ECM content (Fig. 4F, G). Both mutants showed a visible increase in the surface volume of the limbs. In addition, all *TBX3* mutants displayed fusion of at least two digits (Fig. 4H). Transversal sections of these limbs confirmed the retention of the tissue between digits II and III, resembling cutaneous syndactyly in both mutants (Fig. 4I). Quantification analyses of these images revealed a significant increase in the overall autopod volume, cell number and connective tissue content in both mutants (Fig. 4J-L). These results indicate that the expression of *MEIS2* and *TBX3* in the distal and interdigital mesenchyme can recapitulate essential aspects of bat wing development. This includes increased proliferation and matrix production, resulting in retention of interdigital tissue with consecutive fusion of digits. Overall, these analyses support that the distal activation of a gene program mediated by *MEIS2* and *TBX3* plays a role in bat chiropatagium formation.

## Discussion

This study aims at elucidating the molecular basis and cellular origin of the interdigital wing membrane of bats, the chiropatagium. Previous studies have attempted to identify the genes and mechanisms behind this fascinating evolutionary adaptation. Candidate gene approaches, for instance, suggested an involvement of pro-apoptotic factors such as *BMPs* (*42*) and their antagonist *GREM1* (*13*) or a second wave of *SHH* expression in the interdigital space (*43*). More systematic approaches using transcriptional profiling and the integration of regulatory data identified genes of the *HoxD* cluster as well as components of canonical Wnt-signaling (*17*). Yet, these genome-wide studies lacked cellular resolution and therefore much remains elusive. Here, by using scRNA-seq, we were able to assign expression patterns to specific cell populations thereby disentangling previous contradictory observations. Cells expressing pro-apoptotic factors in bats are equivalent to the RA-Id cluster observed in mice, where interdigital regression takes place. In contrast, *GREM1* is expressed in bat fibroblast cells, which originate from a distinct developmental trajectory from the RA-Id cluster and constitute the major component of the chiropatagium (Fig. 2O). Nonetheless, besides apoptosis, other mechanisms including epidermal cell migration (*37*), and the remodeling of ECM components are involved in interdigital tissue regression (*44*). This, together with our results, indicates that apoptosis is not sufficient for sculpting the digits in mammals. Indeed, further analysis of the fibroblast gene program identified an enrichment of genes associated with these processes, i.e. ECM organization, cell migration and proliferation. Alterations in the balance between cell death, and proliferation and migration are likely to change the interdigital cell composition and can result in the retention of interdigital tissue (syndactyly) (*27, 37, 45*).

A major challenge in comparative single-cell analyses lies in data integration, which risks overcorrection and the consequent masking of biological variation (*46*). This was also of concern during our integration of bat and mouse data, where the interdigital distal fibroblasts forming the bat chiropatagium clustered together with other fibroblasts from both species. However, data from micro-dissected chiropatagium revealed that such clustering was not artifactual, but rather reflected the use of similar transcriptional programs. The remarkable similarity we found between cells of such strikingly different limbs raises the question of how this difference is achieved. It is well-documented that during evolution, the same set of genes is often reused through processes like convergent evolution or deep homology conservation (*47, 48*). For example, the formation of lateral patagia enabling gliding has independently appeared multiple times in marsupials through convergent evolution, where the upstream factor *Emx2* is activated by distinct regulatory elements in different glider species (*49*).

Here, we identify a gene program that has been repurposed through evolution, where two TFs, MEIS2 and TBX3, appear among the primary regulators. Specifically, we show that MEIS2 is a potential direct activator of many other TFs in bat wings, regulating other downstream genes. Both factors were previously described in different bat species (*Miniopterus natalensis*, *Miniopterus schreibersii*) as expressed in the distal bat FL (*15, 16*), indicating a conserved function in wing development. *Meis2* has also been previously reported to be expressed in distal E14.5 mouse limbs, based on *in situ* hybridization signal (*14*). However, our quantification based on scRNA-seq demonstrates that the expression levels are low and are present in markedly fewer cells compared to the bat FL (Fig. 2R). In contrast, *Meis1* and *Meis2* (Meis) homeobox TFs are well known to be robustly expressed early (<E11.5) in the proximal part of the limb, where they determine the identity of stylopod and zeugopod vs. autopod (*50*). Accordingly, mutating Meis TFs results in limb shortening due to altering the proximo-distal segmental borders (*34*). The specification of proximal identity by *Meis* genes is an ancient function conserved across the vertebrate lineage, including mammals (*34*), birds (*50, 51*) and amphibians (*52*). Interestingly, in Drosophila, the *Meis* homologue *hth* is also required for proximal leg development (*50*). Likewise, *Tbx3* is a gene expressed in the proximal limb mesenchyme and plays a crucial role during limb patterning in establishing anterior-posterior boundaries (*53*). The importance of *MEIS2* and *TBX3* in chiropatagium development is supported by our studies in transgenic mice. The gene expression changes observed in mutant limbs, together with the alterations in morphology, cell number and matrix production, reflect key features associated with the gene program of chiropatagium cells (Fig. 3). Thus, the ectopic expression of *MEIS2* or *TBX3* in interdigital distal cells induces a gene program that partially resembles that observed in bats and leads to tissue retention. It is likely that the expression pattern of *MEIS2* observed in bats requires regulatory changes rendering *MEIS2* susceptible to specific FL autopod signals, such as a FL specific Hox code (*54, 55*). This may encompass the observed activation of 3’ anterior Hox paralogs like HOXA1/2.

Phenotypic evolutionary innovations can, in principle, arise from gene duplications or losses. Yet, a genomic comparison of six bat reference genomes failed to reveal expansion or loss of any candidate gene that might play a role in limb development (*56*). Alternatively, already existing genes can be newly recruited into regulatory gene networks (co-option) (*57, 58*), or regulatory changes can modify gene expression within existing ones (*49, 59, 60*). Instead, our data point towards a high degree of similarity in gene expression between species suggesting the reuse of a transcriptional program already existent in the limb but at a different anatomical position. It is probable that the re-use of this gene program occurs within a markedly disparate epigenetic landscape, thereby activating slightly disparate and novel gene sets. A similar scenario has recently been reported for skeletogenic cells found in different parts of the body (*61*). Chondrocytes that originate from different germ layers use distinct sets of regulatory elements for activation of similar gene programs. Like the chiropatagium cells, which are equivalent at the transcriptional level to the proximal fibroblasts but diverge at the gene regulatory level. Thus, even a change as drastic as the development of a wing from individual digits can apparently be achieved by relatively small changes and the repurposing of already existing and active pathways. Following the principle of parsimony, evolution constructed novelty by making minimal modifications to already existing elements.

## Methods

### Animal Samples

#### Mice

Wildtype mouse embryonic tissues were derived from crossings of CD1 x CD1 or C57BL/6J x 129. Transgenic embryos were generated by tetraploid aggregation (*1*). Female mice of CD1 genetic background were used as foster mothers. Mice were kept in a controlled environment (12 h light and 12 h dark cycle, temperature of 20–22.2 °C, humidity of 30–50%) and water, food and bedding were changed regularly.

All animal experiments and procedures were conducted as approved by LAGeSo Berlin under the following license numbers: ZHV120, G0176/19-MaS1_Z, G0243/18-SAld1_G and G0098/23-SAld1_G.

#### Bats

Bat samples (*Carollia perspicillata*) were obtained from a captive population maintained at the Papiliorama zoo in Kerzers, Switzerland. To control population growth, some individuals were occasionally culled by cervical dislocation performed by trained personnel, following general guidelines for animal handling and in vivo research (*2*). If pregnant females were present among the culled individuals, embryos were dissected and preserved for different molecular procedures. Females in late pregnancy were not culled for ethical reasons. In addition, bat samples from *Carollia perspicillata* were obtained from a breeding colony housed at the Institute for Cell Biology and Neuroscience, at the Goethe University in Frankfurt am Main (keeping permit authorized by the RP Darmstadt). Samples collected from the Frankfurt colony originated from bats that were euthanized for collecting brain tissue without any further experimental manipulation (following § 4 Abs. 3 of the German TierSchG). In female bats, after euthanizing, we additionally checked for possible pregnancies (undetectable from the outside) and embryos were dissected whenever present.

### Genome Annotation

To generate an annotation of *Carollia perspicillata*, we collected transcriptomic data from long-(IsoSeq) and short-RNA reads, mapped those to a chromosome-scale assembly (kindly provided by the Micheal Hiller Lab), and integrated gene predictions using human (hg38), mouse (mm10) and another phyllostomid bat (*Phyllostomus discolor*) as reference annotations. Briefly, IsoSeq data was first analyzed as in (*3*) to produce high-quality ORF predictions. Then, we implemented a strategy to classify and filter transcripts-based features such as known canonical splice sites, known non-canonical splice sites, novel canonical-splice sites, and novel non-canonical splices. A small set of transcripts, with suboptimal features were not used as input for the gene annotation. For example, fusion transcripts (chimeras that include more than one gene), intra-priming (transcripts with more than 85% or at least 10 contiguous adenines within 20 bp upstream of the 3’ end), low coverage (transcripts supporting coding regions by less than 3 reads), RT-switching predictions (an exon-skipping pattern due to a retrotranscription gap caused by secondary structures in expressed transcripts), nonsense-mediated decay (premature stop codons), and intron retention were features all identified as suboptimal. However, when possible, some of these transcripts were used to annotate UTRs. Transcript features used for classification were identified using TAMA-GO (*4*). Then, new TOGA predictions were generated using an updated version (*5*) (https://github.com/hillerlab/TOGA v. 8f09391). We used as reference genomes human (hg38), mouse (mm10), and the pale spear-nosed bat (*Phyllostomus discolor*). Finally, additional RNAseq data from tissues were generated and analyzed, as described in (*3*). Evidences (RNAseq transcripts, reclassified IsoSeq transcripts, TOGA predictions, and proteins) were integrated using EVM (*6*), and downstream steps to annotate non-overlapping UTRs, enrich the annotation with non-coding RNAs, and assign gene names were performed as described in (*3*).

The sensitive prediction of genes including UTRs led to typical artifacts where a gene name was assigned more than once. This sometimes caused a mislabelling of an orthologous gene compared to a reference genome annotation (here hg19). Additionally, in some instances (e.g. HOX gene clusters) transcripts annotated with a unique coding sequence (CDS) were grouped artificially into a single gene based on shared UTR exons. To correct these issues conservatively we used sequence conservation to human (hg19) determined via a one-to-one comparison of both genomes via the alignment software LAST (*7*) with the following parameters: lastdb - uMAM8 -R11 -c; last-train --revsym --matsym --gapsym -E0.05 -C2; lastal -m10 -E0.05 -C2.

As a prerequisite we lifted human CDS regions to the Carollia genome. In cases where a CDS overlaps a conserved region, but a boundary was not conserved, the boundary was interpolated via its distance to the closest overlapping conserved region (approximate lift-over). Known fusion genes and lifted genes with unusually large intron size > 30kb were excluded from subsequent renaming or boundary adjustments.

A Carollia transcript was reassigned to a reference gene name if at least 50% of the original CDS boundaries matched the lifted coordinates of the reference annotation, and if other transcripts of the original gene share less CDS boundaries to another reference gene. Once transcripts and genes were renamed transcripts extending beyond the boundaries of the orthologous reference gene were clipped at the 5’/3’ UTRs. Once transcripts and genes were renamed transcript extending beyond the boundaries of the orthologous reference gene were clipped at the 5’/3’ UTRs.

Finally, for genomic regions without any gene annotation we transferred exon annotations from hg19 to Carollia via approximate lift-over. While this procedure may detect mainly approximate or partial gene annotations it allowed to recover additional ∼500 genes (e.g. XIST) otherwise excluded from analyses. The genome annotation resulted in 23,315 transcripts from 18,697 genes. To add long non-coding transcripts, StringTie (*8*) was used on the short-read RNA-seq data to obtain a transcriptome annotation which was processed further with PLAR (*9*) to identify lncRNAs. The ones not overlapping the initial transcriptome were added to result in 20,421 additional transcripts from 16,141 additional genes.

### Single cell RNA sequencing

#### Single cell isolation, methanol preservation and rehydration

For single-cell gene expression analysis, mouse and bat embryonic limb tissues were dissected and dissociated with trypsin. Cells were filtered through a 40 µm Flowmi Cell Strainer (Merck, #BAH136800040) and pelleted at 300 x g at 4°C for 5 minutes. Cells were resuspended in 1 volume 0.04% BSA/PBS and dehydrated by slowly adding 9 volumes of 100% methanol. Samples were stored at −80 °C until library preparation.

For rehydration, dehydrated cells were centrifuged at 1000 x g at 4°C for 10 minutes and washed twice in 1 mL of rehydration buffer (1 % BSA, 0.4 U/µL Ambion RNse Inhibitor (Invitrogen, #AM1682) and 0.2 U/µL SUPERaseIn RNase Inhibitor (Invitrogen, #AM2696) in 1x DPBS). After the second wash, cells were resuspended in rehydration buffer, counted and diluted to a concentration of 1,000 cells/µL.

#### 10X Genomics single-cell RNA-sequencing library preparation

Single cell gene expression libraries were prepared using the 10X Genomics Chromium Next GEM Single Cell 3’ GEM, Library & Gel Bead Kit v3.1 (10X Genomics, #PN-1000121) according to the manufacturer’s instructions. The aimed target cell recovery in each experiment was 10,000 cells.

To generate Gel Beads-in-emulsion (GEMs), reactions were assembled in a Chromium Next GEM Chip G (10X Genomics, #PN-1000121). Chips were run on a Chromium Controller X/iX. Sample indices were added to the cDNAs via PCR using the Single Index Kit T Set A (10X Genomics, #PN-1000213) or Dual Index Kit TT Seat A (10X Genomics, #PN-1000215). Library concentration was measured by Qubit dsDNA HS Assay Kit (Invitrogen, #Q33231) and quality was assessed using Bioanalyzer High Sensitivity DNA Analysis Kit (Agilent, #5067-4626). Finally, scRNA-seq libraries were sequenced on an Illumina NovaSeq 6000 with asynchronous 28 bp / 90 bp paired-end reads. Single-cell experiments were performed in biological duplicates.

### Single cell RNA analysis

#### Filtering, normalization and integration

The different single-cell RNA-sequencing libraries were processed using 10X Genomics CellRanger v6.0.2 (*10*) and our custom genome annotations for *C. perspicillata* and *M. musculus*. Individual count matrices were filtered for quality based on relative UMI counts (removed >4*mean and <0.2*median of the sample), percentage of ribosomal UMIs (removed >median + 3*MAD & <median + 3*MAD), and relation of UMI count/genes detected (removed <0.15 & UMI count <2/3). The filtered datasets were integrated in a species / limb manner (e.g. all Mouse FL datasets together). For this we used Seurat v4.3.0 (*11*), first Log normalizing each dataset with a factor of 10’000 and scoring the cell cycle state of each cell. Then, using the SCTransform tool we regressed the percentage of ribosomal UMIs, the UMI count, and the S and G2M cycle score of each cell. Using the top 25% integration features, we found integration anchors using the SCT normalization, 20 first dimensions and a k.filter of 100. These anchors were used with the tool IntegrateData.

#### Dimensionality reduction and clustering

Using the integrated data from the top variable features (stand. variance > median + MAD of the data) we calculated a PCA and used the first 20 (18 for the Chiropatagium samples) PCs downstream. We calculated tSNE plots using FFT-accelerated Interpolation-based t-SNE (*12*) in order to retain and represent the global structure of the data (*13*). Clusters were computed using FindNeighbors and FindCLusters with a resolution of 0.7 and 42 as a random seed. Marker genes for each cluster were then calculated from the un-integrated expression data using a minimum percentage of expression of 0.25, and a LFC threshold of 0.5.

The cells identified as LPM-derived were subsetted, and the same process was followed to generate the presented individual datasets. From the MmHl dataset, a cluster with a high proportion of Haemoglobin UMIs was removed.

#### Interspecies atlas

The integration of all Cp and Mm limb LPM-derived cells followed the same logic. We subsetted the LPM cells from each of the datasets, separated the individual libraries, and used the same workflow. For this integration we used the top 12.5% of the integration features. To find clusters we used a resolution of 0.6. The identity labels of this integration were used for the individually clustered datasets (eg. Mm FI). Each cluster was first classified as one of the three main groups of LPM cells by simple majority, and then was given the identity of the most represented label from that group.

#### Apoptosis-related expression comparison

The cells from the integrated cluster 3 RA-Id were compared against the rest of the cells in a species / limb fashion (eg. cells within the Mm Fl dataset) using the function FindMarkers with a minimum expression percentage and a LFC threshold of 0.0001, using all genes related to apoptosis, the marker genes from the cluster, and 20 random marker genes from other clusters.

#### Label transfer to Chiropatagium cells

To transfer annotation labels from the CpFl-LPM dataset to the Chiropatagium-LPM cells, we found transfer anchors using the first 20 PCs and the SCT normalized data and used the TranferData function.

#### Pseudotime analyses

We calculated RNA velocities using velocyto (*14*) on our single-cell libraries, using the stricter mode for the Cp samples. From each individual LPM sample, we subsampled the Hoxd13+ cells (>0 UMIs). We re-clustered and annotated the dataset, and then exported it to an AnnData format and integrated the RNA velocity to be further analyzed. Using scvelo v0.3.2 (*15*) we filtered the data and found the first and second order moments with the 20 first Pcs. We then ran the dynamical model and calculated RNA velocity allowing for differential kinetics. Guided by the apparent RNA velocity trajectories, and based on the identities of the clusters, we subsetted the data further to remove the chondrogenic lineage as much as possible. We then generated diffusion maps using the first 15 PCs and chose the diffusion eigenvectors 1 and 2. Using the slingshot package we inferred trajectories of differentiation and pseudotime values for each cell using the seurat-calculated clusters, and the apparent end and start clusters according to the RNA velocity. With this data, we again computed RNA velocity without a dynamical model. Using CellRank v2.0.4 (*16*), we computed velocity, connectivity and pseudotime kernels which were combined in proportions of 0.2, 0.4, and 0.4, respectively.

#### Distal - Proximal computational dissection

Each species / limb dataset was given a proximal or distal score using Seurat. With the function AddModuleScore we scored each cell for the sets of genes distal (*MSX1*, *HOXD13*), proximal (*SHOX2*), Chondrogenic (*SOX9*, *COL2A1*), and fibroblastic (*DCN*, *ZFHX3*). The same approach was used for other sets of genes. Per cluster, we calculated the mean of the difference between the proximal - distal scores. We then assigned clusters with the ⅓ most extreme score differences as very distal / proximal. We then categorized genes as typical proximal or distal if in both species they are expressed in at least 20% of the cells of 20% of the corresponding clusters, and they show a difference of > 0.15 LFC against the Opposite cells. Genes highly expressed in the chondrogenic lineage were excluded. In Supplementary Fig. 4a and b, we show the top (by LFC) 15 genes expressed in < 15% of the opposite cells. Co-expression of genes is measured as UMIs_x_ > 0 & UMIs_y_ > 0.

#### Proximo / Distal Fibroblast expression program

In order to understand what defined the expression profile of proximal and distal fibroblasts, without detecting the differences between them, the distal fibroblasts (10 FbI1) were compared against the rest of the cells, except proximal fibroblasts (cells from 8 FbA and 9 FbL labelled as 9 FbL in the main integration), and vice-versa. The fibroblast program 2 was done in the same way with cells from cluster 7 FbIr and cells from 8 FbA labelled as 8 FbA in the main integration. This was done in two rounds, first using the highly variable genes, and then using all the genes that had been detected as DE in both comparisons. GO Terms enrichment analyses were made using clusterProfiler (*17*) and all the genes expressed in at least 9 cells in the sample as the background universe. The function simplify was then used with a cut-off of 0.6 on the adjusted p value.

#### Mutants’ analysis and differential expression

The single - cell datasets from mutant mice were subsetted for LPM cells expressing Hoxd13 and integrated with corresponding cells from the mouse wild type forelimb. Each of the new clusters found was annotated on the wt Mm Fl dataset following the logic before. Using MAST (*18*), Cluster- wise we tested for differential gene expression between the wt and mutant cells. For this we considered the highly variable genes, all the genes part of our proximal / distal fibroblast program, and excluded all genes on the x chromosome, mt and ribosomal genes. Only those genes expressed in at least 15% of the cells from either genotype were tested using MAST using a zlm with the formula “genotype + orig.ident + percent.rp”. We then calculated an lrTest on the genotype coefficient and a subsequent p value adjustment using p.adjust. A hypergeometric test was used to assess the over representation of the DE genes (p.val < 0.01 LFC > 0.25) in our program. GO Terms were calculated on the totality of genes overexpressed in the mutant cells per cluster using the approach described above. We filtered duplicated terms based on the set of genes present. We then manually grouped the top 10 terms by adjusted p value of all clusters in the categories presented in Fig. 4d. We calculated the mean expression of the genes in our proximal / distal fibroblast program in the cluster 10 FbI1 of the but FL data, and then calculated the Pearson correlation of each cell to this mean using the same genes. For this, we focused on the clusters where we found Over expression of Meis2 and Tbx3 (p.val < 0.01 LFC > 0.15) in the mutants and the corresponding clusters in the Mm Fl and Cp Fl data sets.

### Fluorescent microscopy apoptosis assays using LysoTracker and Immunofluorescence against Cleaved Caspase-3

Bat embryonic limbs were dissected in cold DPBS and separately processed for the two different cell death assays.

For the lysosomal staining, samples were transferred immediately into 5 µM LysoTracker Red DND-99 (Invitrogen, #12090146) in DPBS and incubated at 37 °C for 45 minutes, then washed four times in DPBS and fixed overnight in 4% PFA/DPBS. After that, samples were washed for 10 minutes in DPBS, dehydrated through a methanol series (25, 50, 75 and 100%) and stored at −20 °C until imaging.

For the caspase assay, samples were fixed in 4% PFA/DPBS for 1 to 2 hours at 4 °C, then washed three times and stored in DPBS at 4 °C until immunofluorescent staining was performed. Samples were washed twice in DPBS for 5 minutes, then permeabilized in 0.5% Triton-X-PBS (PBST) (3 x 1 hour incubation) and blocked in 5% FCS/PBST overnight at 4 °C. Anti-Cleaved-Caspase-3 (D175) Antibody (rabbit polyclonal, Cell Signaling Technology, #9661, Lot 47) was diluted in blocking solution (1:400) and incubated for 72h at 4 °C. Samples were washed three times with blocking solution and three times with PBST, then incubated in blocking solution overnight at 4 °C. Donkey anti-Rabbit IgG (H+L) Highly Cross-Adsorbed Secondary Antibody, Alexa Fluor™ 568 (Invitrogen, #A10042, Lot 2306809) and DAPI were diluted in blocking solution (1:1000) and incubated for 48h at 4 °C. Samples were washed three times with blocking solution, three times with PBST, three times with DPBS, and post-fixed in 4% PFA/DPBS for 20 minutes.

#### Confocal Fluorescence Imaging

At this point, samples from both experiments were similarly treated. They were washed three times with 0.02M phosphate buffer (pH 7.4) and cleared in RIMS (Refractive Index Matching Solution, 13% Histodenz (Sigma-Aldrich D2158) in 0.02M PB) at 4 °C for at least one day. Whole-mount limbs were then imaged with a Zeiss LSM880 confocal laser-scanning microscope in fast-Airyscan mode. Images were then processed with the ZEN software (Airyscan processing and Max intensity projection of the Z-stacks) and with Fiji.

### Gene regulatory network analysis

Gene regulatory networks were generated using the Python implementation of SCENIC (pySCENIC) (*19*).

Raw counts, without SCT-transform, and cell-type identities were extracted from the generated Seurat object. These counts were then filtered as described by pySCENIC authors. This included, filtering out cells with less than 200 or more than 6000 genes with counts, and filtering genes appearing in less than 3 cells.

Vertebrate motifs were downloaded from JASPAR at the following link: https://jaspar.elixir.no/download/data/2024/CORE/JASPAR2024_CORE_vertebrates_non-redundant_pfms_jaspar.zip. These motifs were converted to clusterbuster motifs using Biopython’s motif submodule, for input into pySCENIC. Transcription factor and motif names were also extracted from the downloaded motifs.

Adjacencies between these transcription factors were calculated with the filtered counts, using pySCENIC’s GRN function. Regulons, highlighting enriched motifs, were then calculated from these adjacencies using pySCENIC’s CTX function. Cells where the transcription factor or target gene expression were 0 were masked when calculating correlation between a transcription factor-target pair, both positive and negative regulons were calculated, and no pruning was performed. pySCENIC’s default behaviour is to prune regulons based on cisRegulatory information, however due to lack of compatibility with the novel bat annotation, this step was skipped. Hence, all enriched motifs are included, and regulons were filtered downstream.

The area under the curve (AUC) was then calculated on these regulons using the filtered counts, to determine regulon enrichment using pySCENIC’s AUCELL function. This AUC information was combined with cell-type labels to generate regulon specificity scores (RSS) for both positive and negative regulons across all cell-types.

Finally, these RSSs were used to generate gene regulatory networks, determining the edges between regulon nodes. Only transcription factor-target gene connections with a weight (adjacency) greater than 10 were included, filtering out weak transcription factor-target pairings. Moreover, target genes with a mean expression across all cells less than 0.05 were also excluded.

### RNA sequencing and analysis

Total RNA was extracted using RNeasy Mini Kit (Quiagen, #74106) or RNeasy Micro Kit (Quiagen, #74004) according to the manufacturer’s instructions. Limb samples derived from E11.5 and E13.5 as well as CS15 and CS17 embryos were directly homogenized in RTL buffer supplemented with 1% β-Mercaptoethanol and applied to spin columns. Limb tissues from older embryonic stages (E15.5 and CS19) were crushed in liquid nitrogen using Bel-Art SP Scienceware Liquid Nitrogen-Cooled Mini Mortar prior to homogenization. Genomic DNA was removed using the RNase-Free DNase Set (Quiagen, 79254).

For gene expression analysis, samples were poly-A enriched and libraries were prepared using the Kapa HyperPrep Kit (Roche, #07962347001). RNA libraries were sequenced on a Novaseq2 with 100 bp paired-end reads. RNA-seq experiments were performed at least in biological duplicates. Read mapping to mm39 and *Carollia perspicillata* reference genomes was performed using the STAR_2.6.1d software (*20*) with the following options: -- chimSegmentMin 10 --alignIntronMin 20 --outFilterMismatchNoverReadLmax 0.05 -- outSAMmode NoQS --outFilterMismatchNmax 10. For samples obtained from transgenic animals the transgene sequence was temporarily merged with the genome annotation using the option -add. For each sample, read counts per gene were obtained via the R function “featureCounts”, with the parameter --countReadPairs -s 2. For visualization, CPM normalized bigwig files were created using the “bamCoverage” tool from deepTools (*21*) or read counts were normalized to reads per kilobase million (RPKM) based on the number of uniquely mapped reads.

#### Differentially expressed genes

Differentially expressed genes (DEGs) were identified from featureCounts (*22*) count matrices using DESeq2 v1.38.3 (R v4.2.2) (*23*). For each comparison, replicate quality was assessed using PCA and Euclidean distance between samples. When necessary, outlier replicates were removed from further analysis. Lowly expressed genes were then filtered using edgeR v3.40.2 filterbyexpr (*24*) and non-annotated transcripts were removed to ensure 1-to-1 comparisons between the species. Counts were normalized and differential expression calculated as log2 fold change (LFC) from the mean of normalized counts. Genes were assigned as differentially expressed for LFC larger than ±0.5 and adjusted Wald test p value below 0.05.

### ChIP sequencing and analysis

Mouse and bat embryonic limbs were dissected and fixed in 1% formaldehyde in 10% FCS/PBS and subsequently snap-frozen and stored at −80°C until further processing. Chromatin immunoprecipitations were performed using the iDeal ChIP-seq Kit for Histones (Diagenode, #C01010051) and iDeal ChIP-seq Kit for Transcription Factors (Diagenode, #C01010055) according to the manufacturer’s instructions. Briefly, fixed limbs were lysed and sonicated using a Bioruptor Plus Sonication device (45 cycles, 30s on, 30s off, at high power setting) in provided buffers. 5 µg sheared chromatin were used for histone immunoprecipitation with 1µg of the following antibodies: anti-H3K4me3 (Millipore, #07-473), anti-H3K4me1 (Diagenode, #C15410037), anti-H3K27ac (Diagenode, #C15410174). For Meis2 immunoprecipitation, 20 µg of sheared chromatin were used with two anti-Meis antibodies simultaneously, one recognizing the conserved C-terminal domain of Meis1a and Meis2a and the other recognizing all Meis2 isoforms as previously described (*25*). 2µg of each antibody were used per immunoprecipitation. Antibodies were generously provided by Dr. Miguel Torres. Libraries were prepared using the Kapa HyperPrep Kit (Roche, #07962347001) and libraries were sequenced on a NovaSeq2 with 100bp paired-end reads. ChIP-seq experiments were performed in biological duplicates.

E10.5 Meis1/2 ChIP data from mouse forelimbs was obtained from previously published data (*25*).

Read mapping of the sequenced samples to mouse and bat reference genomes (mm39 / carPer2) was performed using the STAR_2.6.1d software (*20*). Reads were then filtered and sorted and duplicates were removed using SAMtools (*26*). For visualization, CPM normalized bigwig files were created using deepTools “bamCoverage” tool. Input samples were subtracted using deepTools bamCompare (*21*).

#### H3K27ac differential peak and motif enrichment analysis

Differential acetylation regions between dFL and dHL, as well as acetylation regions common in both conditions, were predicted from ChIP-seq alignments using macs2 (*27*) bdgdiff command with the following parameter: -l 800 -g 500 and a likelihood ratio cut-off of 1000.

The coverage of dFL-specific acetylation regions relative to the acetylation regions shared between dFL/dHL was calculated for each TAD.

The dFL-specific acetylation regions, and the acetylation regions shared between dFL/dHL were first intersected with accessible regions in limbs (CS17). The dFL-specific accessible acetylation regions were given as input and the commonly accessible acetylation regions as background for motif enrichment analysis done by homer2 (*28*). A q-value cut-off of 0.01 was used for the enriched motifs. The list of significantly enriched motifs was manually curated so that only the most significant TF motifs in each gene family were retained.

Tornado plots were generated for peak distribution visualization using deeptools v3.5.4. (*21*) The scores per regions were calculated using computeMatrix in scale-regions mode, where the scores were based on the ChIP bigwig pileup files and the regions based on BED files defining the ChIP peaks.

#### MEIS binding analysis

Meis2 binding signal of one of our dFL replicates was aggregated by TADs. The signal (area under the curve) was calculated using deepTools pyBigwig (*21*). By analyzing the second derivative of the density distribution of the signal by TAD, we found the dividing point to the subpopulation of enriched TADs. The same procedure was carried out for the coverage length of the acetylation peaks. Genes were categorized as “Cytoskeleton”, “ECM”, or “Transcription Factor” using the Uniprot database, if the Keywords included the terms “Cytoskeleton”, “Extracellular matrix”, or (“DNA-binding” OR “Transcription regulation”) respectively.

### ATAC sequencing

ATAC-seq protocol and library preparation was performed as previously described (*29*). In short, bat limbs were dissected and dissociated with trypsin. 50,000 cells per reaction were lysed and isolated nuclei were incubated with Tn5 Transposase (Illumina, #20034197) for transposition. DNA fragments were purified and barcoded adaptors were added via PCR. Fragments were purified and sequenced as 100 bp paired-end reads on a NovaSeq 6000 system. ATAC-seq experiments were performed in biological duplicates.

For processing, adaptors were trimmed with the cutadapt tool (*30*) and reads were mapped to indexed reference genomes (mm39/carPer2) using Bowtie2 (*31*). Reads were then filtered, sorted and duplicates removed according to ChIP-seq processing. Reproducible peaks were called using Genrich with default parameters.

### *Carollia perspicillata* embryonic fibroblast culture

Head- and organ-free tissue from a CS16/CS17 bat embryo (*Carollia perspicillata*) was minced in DMEM supplemented with 15% FCS and cryo-frozen in DMEM containing 10% DMSO and 15% FCS until further processing. To establish bat embryonic fibroblast culture, tissue pieces were thawed and digested with trypsin for 20 minutes at 37°C. Cells were centrifuged at 1000 rcf for 5 minutes, resuspended in fibroblast culture media (DMEM high glucose, 15% FCS, 1% Pen/Strep, 1% L-glutamine) and transferred to a 6 well plate for cell attachment and expansion. Cells were split into a new culture flask when they reached a density of approximately 80% or cryofrozen in freezing media (DMEM high glucose supplemented with 15% FCS and 10% DMSO) in 1-3x106 aliquots. Fibroblast cells were cultured at 37°C and 5% CO2.

### Hi-C

Hi-C libraries from *Carollia perspicillata* embryonic fibroblast cells or bat embryonic forelimbs were prepared as previously described (*32*). In short, approximately 1×10^6 cells fibroblast and 500,000 limb cells were fixed in 2% formaldehyde in 10%FCS/PBS. After cell lysis, chromatin was digested with *DpnII* enzyme (NEB, #R0543), digested ends were marked with biotin-14-dATP (Invitrogen, #19524016) and subsequently ligated. Crosslinking was reversed, DNA precipitated and sheared to a fragment size of 300-600 bp using a S-Series 220 Covaris sonicator. The biotin-containing fragments were pulled down with Dynabeads MyOne Streptavidin T1 beads (Invitrogen, #65602) and ends were repaired using Klenow Fragment DNA polymerase I (NEB, #M0210) and T4 DNA polymerase (NEB, #M0203). Adaptors were added to DNA fragments using NEBNext Multiplex Oligos for Illumina kit (NEB, # E7335S) and sequencing indices were added by PCR using NEBNext Ultra II Q5Master Mix (NEB, #M0544). Hi-C libraries were generated as three technical replicates and sequenced on a NovaSeq2 as 100bp paired-end reads.

For read processing, the *Carollia perspicillata* reference genome (carPer2) was indexed with the short read aligner BWA 0.7.17 (*33*). Raw reads from sequenced Hi-C libraries were then processed using the Juicer pipeline v1.5.6 (*34*). The three replicates were processed independently and subsequently merged after filtering and deduplication. Hi-C maps with various bin sizes were generated using Juicer tools 1.11.09 (*34*) using the parameter pre -q 30. For displaying Hi-C maps as heatmaps, KR normalized maps with 5 kb bin size were used.

#### TAD calling

Topologically associated domains (TADs) were called using Hi-C data of *Carollia perspicillata* embryonic fibroblasts with the software TopDom (*35*) (KR normalized, resolution: 50 kb, window size: 10).

### Cloning ectopic expression constructs

sgRNA construct targeting the safe harbour locus H11 locus was generated using the same sequence previously described (*36*). sgRNA oligos were cloned into *BbsI* digested and dephosphorylated pSpCas9(BB)-2A-Puro (PX459) V2.0 vector (Addgene; #62988). sgRNA sequences can be found in Table 1.

For cloning of expression constructs, a pUC-Amp plasmid containing homology arms (0.7 kb) designed on the H11 knock-in site was ordered from Twist Bioscience, and the Hsp68 promoter was used as minimal promoter. A previously described interdigital enhancer from the Bmp2 locus 38 was amplified from wild type G4 cell DNA; *Carollia perspicillata MEIS2* cDNA was ordered from Twist Bioscience as a fragment; *Carollia perspicillata TBX3* cDNA was ordered from GeneWiz (Azenta Life Sciences) as a pUC-Amp vector. Kozak sequence (GAGTGG), SV40 polyA signal were included in the design of both overexpression constructs. Backbones and fragments were amplified by PCR (PrimeSTAR GXL Polymerase (Takara, #R050A)), introducing also overlapping sequences necessary for Gibson assembly. Fragments were assembled using Gibson Assembly Master Mix (NEB, #M5510) and cloned into 5-alpha Competent E. coli (NEB, #C2987). Products were validated via restriction digestion and subsequent sequencing. Plasmids were purified using Nucleobond Xtra Midi EF kit (Macherey-Nagel, #740420) before transfection.

For alignment of Meis2 and Tbx3 protein sequences, bat and mouse coding sequences were translated into amino acid sequences using ExPASy (*37*). Sequences were aligned using the Multiple Sequence Alignment tool MultAlin (*38*).

### Mouse embryonic stem cell (mESC) culture

Mouse G4 ES cell culture (XY, 129S6/SvEvTac x C57BL/6Ncr F1 hybrid) was performed as previously described (*39, 40*). Briefly, mouse G4 ESCs were grown on a monolayer of mitomycin-inactivated CD1 mouse embryonic fibroblast feeders on gelatin coated dishes at 37°C and 7.5% CO2. ESC culture medium containing knockout DMEM with 4,5 mg/ml glucose and sodium pyruvate (Gibco, #10829-018) supplemented with 15% FCS (PANSera ES, #P30-2600), 10 mM Glutamine (Lonza, #BE17-605E), 1x penicillin/streptomycin (Lonza, #DE17-603) 1x non-essential amino acids (Gibco, #11140-35), 1x nucleosides (Chemicon, #ES-0008D), 0.1 mM beta-Mercaptoethanol (Gibco, #3150-010) and 1000 U/ml Leukemia Inhibitory Factor (LIF, Chemicon, #ESG1107) was changed daily. mESCs were split every 2-3 days or were frozen at a density of 1x 106 cells/cryovial in ESC medium containing 20% FCS and 10% DMSO and stored in liquid nitrogen.

### CRISPR-mediated genome editing / Knock-In

CRISPR-mediated genome editing was subsequently performed as described previously (*41*). In short, 300,000 G4 ESCs were seeded on CD1 feeders 16 h prior to transfection. For site-specific knock-ins at H11 locus, ESCs were co-transfected with 8 μg of the sgRNA and 4 μg of the KI homology construct. After 24 h, transfected cells were split onto puromycin-resistant DR4 feeders in a ratio of 1:3. For antibiotic selection, cells were treated with puromycin for 48 h. For recovery, mESCs were grown for 4-6 days, after which single colonies were picked into 96-well plates containing CD1 feeders. Cells were grown and split into triplicates after reaching sufficient size. One plate was used for DNA harvesting and genotyping of clones. The other two plates were frozen and stored at −80°C for expansion of positive clones.

Clones were screened for expression construct knockins by PCR detecting site-specific insertion breakpoints. Copy numbers of the insertions were then assessed by qPCR. Positive clones were selected for tetraploid complementation. All cell lines and genotyping primers can be found in Table 1.

### Generation of mutant embryos by tetraploid aggregation

For generation of transgenic embryos, selected mutant ESCs were seeded on CD1 feeders, grown for 2 days and then subjected to aggregation by tetraploid morula complementation, as previously described (*1*). Female mice of CD1 strain were used as foster mothers.

### Whole mount *in situ* hybridisation

MEIS2 transgene mRNAs were detected in embryos by WISH using digoxigenin-labelled antisense RNA probes prepared with DIG RNA labelling mix (Roche, #11277073910). Embryos were dissected, fixed in 4% PFA/PBS overnight, and dehydrated in a methanol series (25%, 50%, 75% methanol in 0.1% Tween in DPBS) on ice. They were stored in 100% methanol at −20°C. For staining, embryos were rehydrated in a reversed methanol/PBST series, bleached in 6% H2O2/PBST for 1 hour, treated with 10 μg/ml proteinase K in PBST for 5 minutes, and re-fixed in 4% PFA/PBS with 0.2% glutaraldehyde and 0.1% Tween 20. After washing in PBST, embryos were incubated in L1 buffer (50% deionized formamide, 5x SSC, 1% SDS, 0.1% Tween 20) at 68°C for 10 minutes, followed by hybridization buffer 1 (L1 with 0.1% tRNA and 0.05% heparin) for 2 hours, and then in hybridization buffer 2 (hybridization buffer 1 plus 1.5 µg digoxigenin-labelled RNA probe per embryo) overnight at 68°C. The next day, unbound probe was washed away using L1, L2 (50% deionized formamide, 2x SSC, 0.1% Tween 20), and L3 buffer (2x SSC, 0.1% Tween 20) for three 30-minute intervals at 68°C. Embryos were treated with RNase solution (0.1 M NaCl, 0.01 M Tris pH 7.5, 0.2% Tween 20, 100 μg/ml RNaseA) for 1 hour, washed in PBST, and blocked in TBST 1 (2% FBS, 0.2% BSA) for 2 hours at room temperature. They were then incubated with 1:5000 anti-digoxigenin-conjugated to alkaline phosphatase antibody (Roche, #11093274910) in blocking solution overnight at 4°C. Unbound antibody was washed off with TBST 2 (TBST with 0.1% Tween 20, 0.05% levamisole). For staining, embryos were washed in alkaline phosphatase buffer (0.02 M NaCl, 0.05 M MgCl2, 0.1% Tween 20, 0.1 M Tris-HCl, 0.05% levamisole), then stained with BM Purple AP Substrate (Roche, #11442074). Finally, stained embryos were imaged using a ZEISS SteREO Discovery.V12 microscope with a Leica DFC420 camera. The primers used to generate the transgene MEIS2 probe can be found in Table 1.

### Limb 3D Imaging

PFA-fixed mouse embryo limb specimens were incubated in a solution of 25 µM DraQ5™, dissolved in PermBlock solution (1% BSA, 0.5% Tween® 20 in PBS), for 12 hours. Following three washes with PBS-T, the stained specimens were dehydrated in increasing methanol concentrations (50%, 70%, 95%, and 99% (v/v) methanol in ddH2O). Subsequently, the specimens were stained in a solution of 1.5 µM eosin Y, dissolved in a 1:1 methanol:BABB (benzyl alcohol:benzoate, ratio 1:2) solution for 4 hours, followed by an optical clearing procedure with BABB solution twice for 4 hours each.

After fluorescence whole-mount staining, optically cleared embryo limb biopsies were imaged using the Lightsheet 7 (Zeiss, Oberkochen, Germany). The stacks were captured with a step size of 2.5 µm and at 5x magnification. The ZEN 3.1 (black edition) software was utilized for the operation of the light sheet microscope and the acquisition of the images. The digital three-dimensional reconstruction of light sheet image stacks was conducted using the IMARIS Microscopy Image Analysis Software (Oxford Instruments, Abingdon, UK).

All quantifications were performed using IMARIS Microscopy Image Analysis Software. For the measurement of the volume of the mouse limb, the autopod region devoid of fingers was considered and analyzed using the Volume function in IMARIS for each condition and n=4 independent specimens. The total number of cells was quantified within DraQ5™-positive nuclei of the autopod, excluding the fingers of the limb, using the Spots function in IMARIS for each condition and from n=4 independent samples. For the quantification of connective tissue in the limbs, Eosin Y-positive structures of the limbs were analyzed using the Volume function in IMARIS for each condition and n=4 independent specimens. The differences of the mean were calculated using a Dunnett test following a one-way ANOVA.

## Data availability

Raw and processed data produced in this work have been deposited in the Gene Expression Omnibus (GEO) repositories: GSE275848, GSE275851, GSE275853, GSE275854, GSE275855.

## Acknowledgements

We thank Asita Stiege and Ute Fisher for their technical support. We thank the Papiliorama Zoo directorate and staff members for their support during sampling. We thank the people from the transgenic and sequencing units from the Max Planck Institute for Molecular Genetics for their assistance. We thank Miguel Torres for kindly providing us with the Meis antibodies. We thank all the members from Mundlos, Lupiáñez and Real groups for their fruitful discussions.

## Author contributions

S.M., F.M.R. conceived the study. S.M., D.L., F.M.R. designed and supervised the experiments. M.S., S.A., A.R.R., G.A., N.F., F.M.R. collected samples in Kerzers. M.S., S.A., J.H., B.K., S.M., F.M.R. collected samples in Frankfurt. M.S. generated all the omics data and performed read mapping. C.F. performed all the single-cell analyses and integrated epigenomic analyses. A.M. produced the genome annotation and S.H. and I.U. further improved it. T.Z. established the genome browser and performed preliminary bulk analyses. S.A., J.G. performed the cell death assays. A.B., M.M.R. performed the gene regulatory network analyses. B.L. analyzed the regulatory domains and bulk epigenetic data. A.A.M. analyzed the bulk RNA data. M.S., S.A. generated the mutant mice. R.Y.B., R.H. performed the 3D imaging of the mutant limbs. R.H., S.H., M.V., M.M.R, M.H., D.L., S.M., F.M.R. provided supervision. M.S., C.F., S.M., F.M.R. wrote the manuscript with the help of all other co-authors.

## Supplementary figures

**Supplementary Figure 1. Related to Fig. 1.**
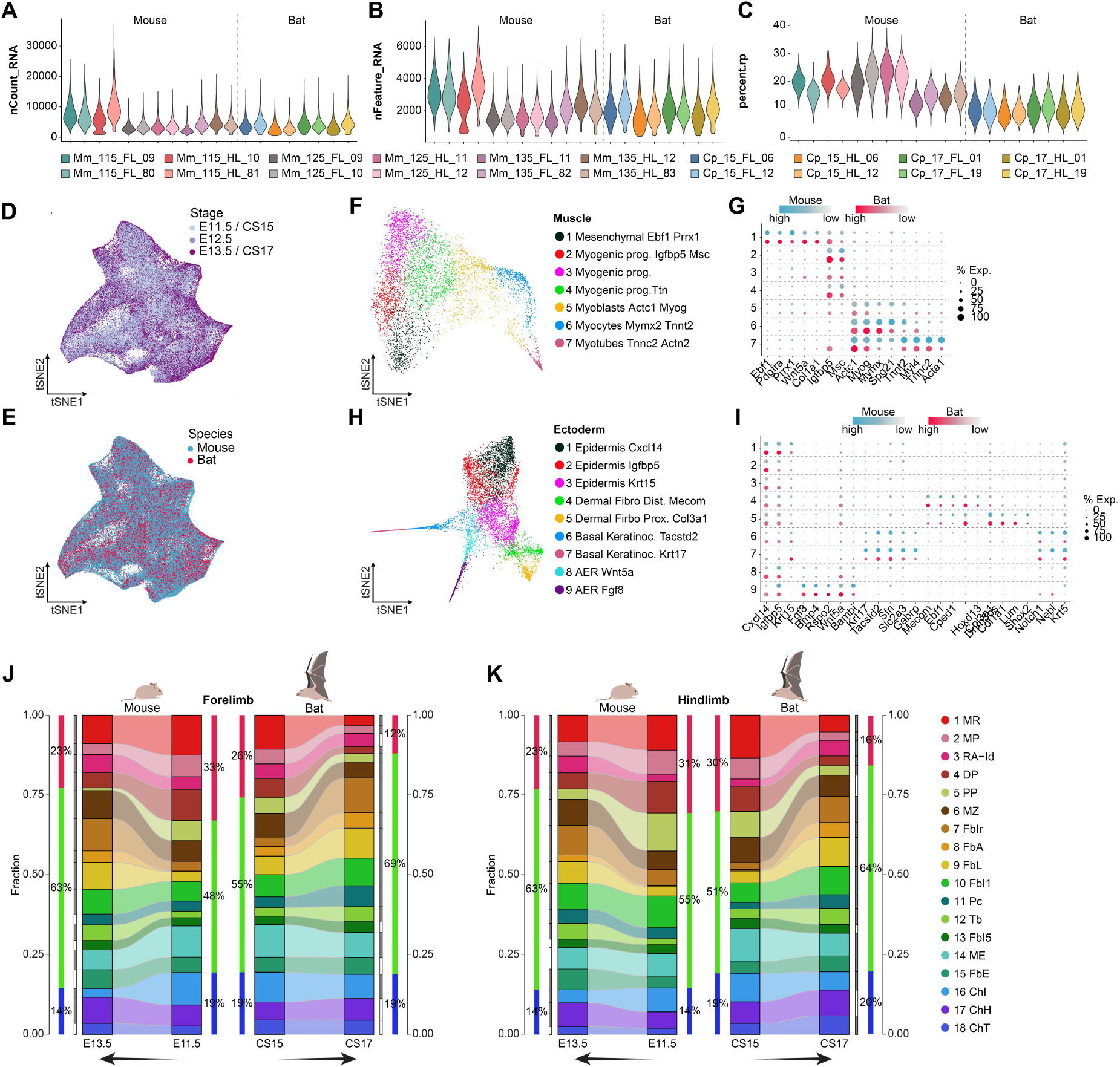
Integrated multi-species single-cell atlas. **A**, **B**, **C** Quality control measurements of all single-cell libraries. **D** Cell contribution of each stage to the integrated atlas. **E** Species contribution to the integrated atlas. **F** Sub-clustering of the muscle cells. **G** Dot-plot showing marker gene expression used for integrated cluster annotation. The color intensity indicates the expression level (blue: mouse; red: bat); the dot size represents the percentage of cells expressing respective marker genes. **H** Sub-clustering of the ectodermal cells. **I** Dot-plot showing marker gene expression used for integrated cluster annotation. **J** and **K** Relative cell proportions over time in mouse and bat forelimbs (J) and hindlimbs (K). Colored side bars represent the main developmental lineages of the LPM-derived cells. Dark gray bars represent significant changes in proportion.

**Supplementary Figure 2. Related to Fig. 1.**
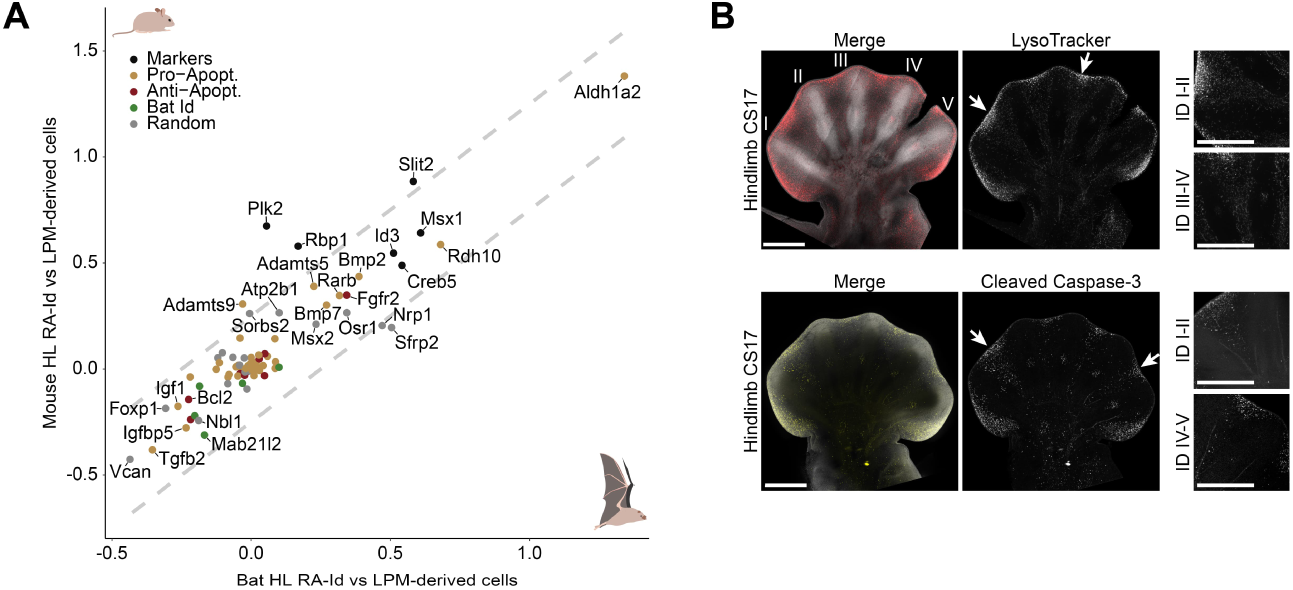
Interdigital cell death in hindlimbs. **A** Correlation of pro- (yellow) and anti-apoptotic (red) genes in the RA-Id cell population of mouse and bat. Marker genes of this cell population are highlighted in blue; genes previously reported to be expressed in bat interdigital regions are highlighted in green. Shown is the lfc of differential gene expression analysis between the RA-Id cluster 3 versus the rest of the LPM-derived mesenchyme per species in the HL. A set of random genes was included as control. **B** Lysotracker staining (upper panel) and immunostaining against Cleaved Caspase 3 protein (lower panel) of bat HL at stage CS17 with magnification of interdigital regions between digits I and II and IV and V (indicated by arrows) shown on right. Merged images show DAPI and LysoTracker or Cleaved Caspase-3 signal. Scale bars represent 500 µm.

**Supplementary Figure 3. Related to Fig. 2.**
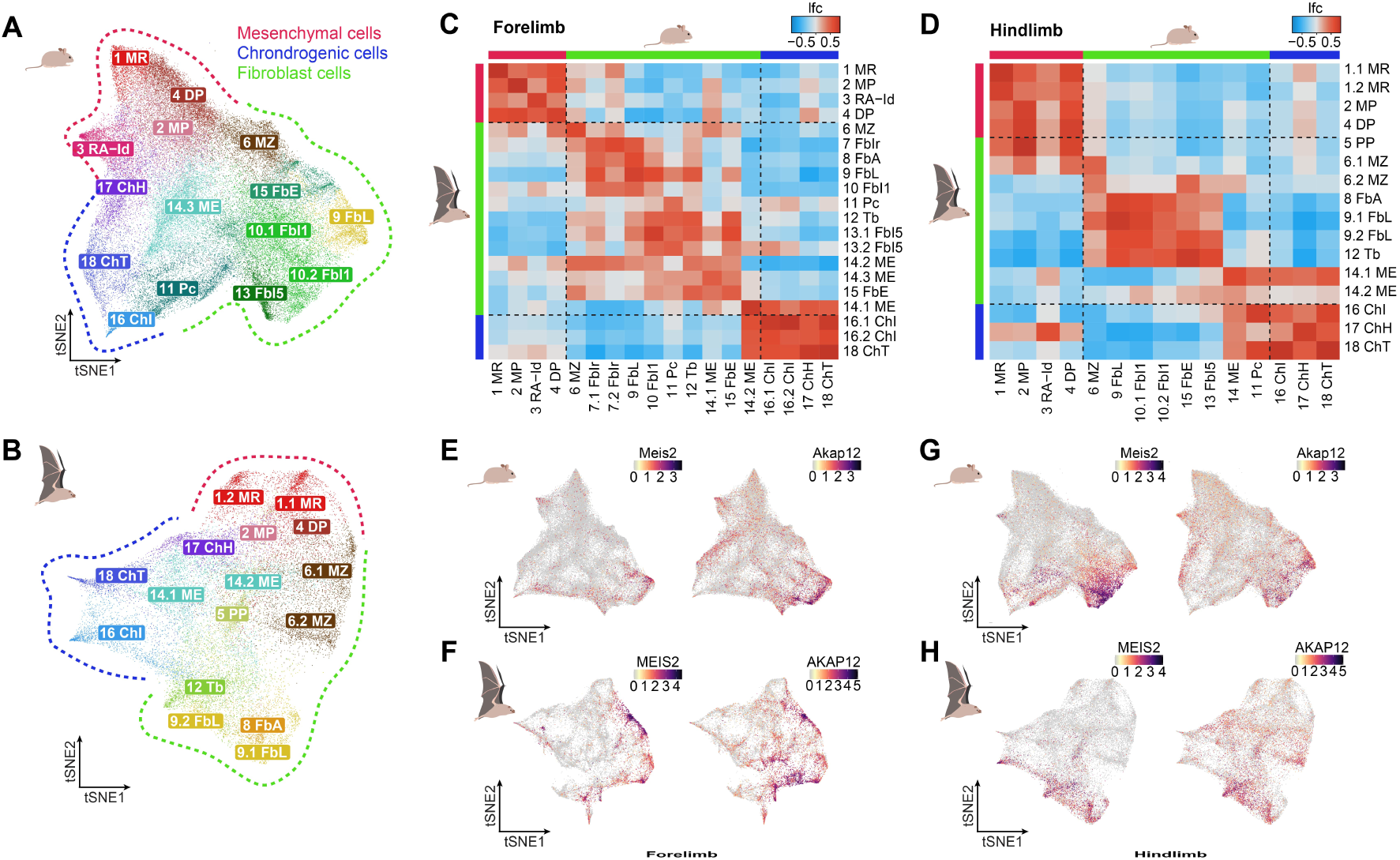
Species-specific clustering and cluster correspondence between species. **A** Individual mouse HL LPM clustering. **B** Individual bat HL LPM clustering. **C** Clustering correspondence between mouse and bat FLs. Shown is the Pearson correlation of the lfc of the top 10 marker genes of all clusters between the focus cluster and all other cells. **D** Clustering correspondence between mouse and bat HLs. **E** Expression pattern of *Meis2* and *Akap12* in the LPM of mouse FLs. **F** Expression pattern of *MEIS2* and *AKAP12* in the LPM of bat FLs. **G** Expression pattern of *Meis2* and *Akap12* in the LPM of mouse HLs. **H** Expression pattern of *MEIS2* and *AKAP12* in the LPM of bat HLs.

**Supplementary Figure 4. Related to Fig. 2.**
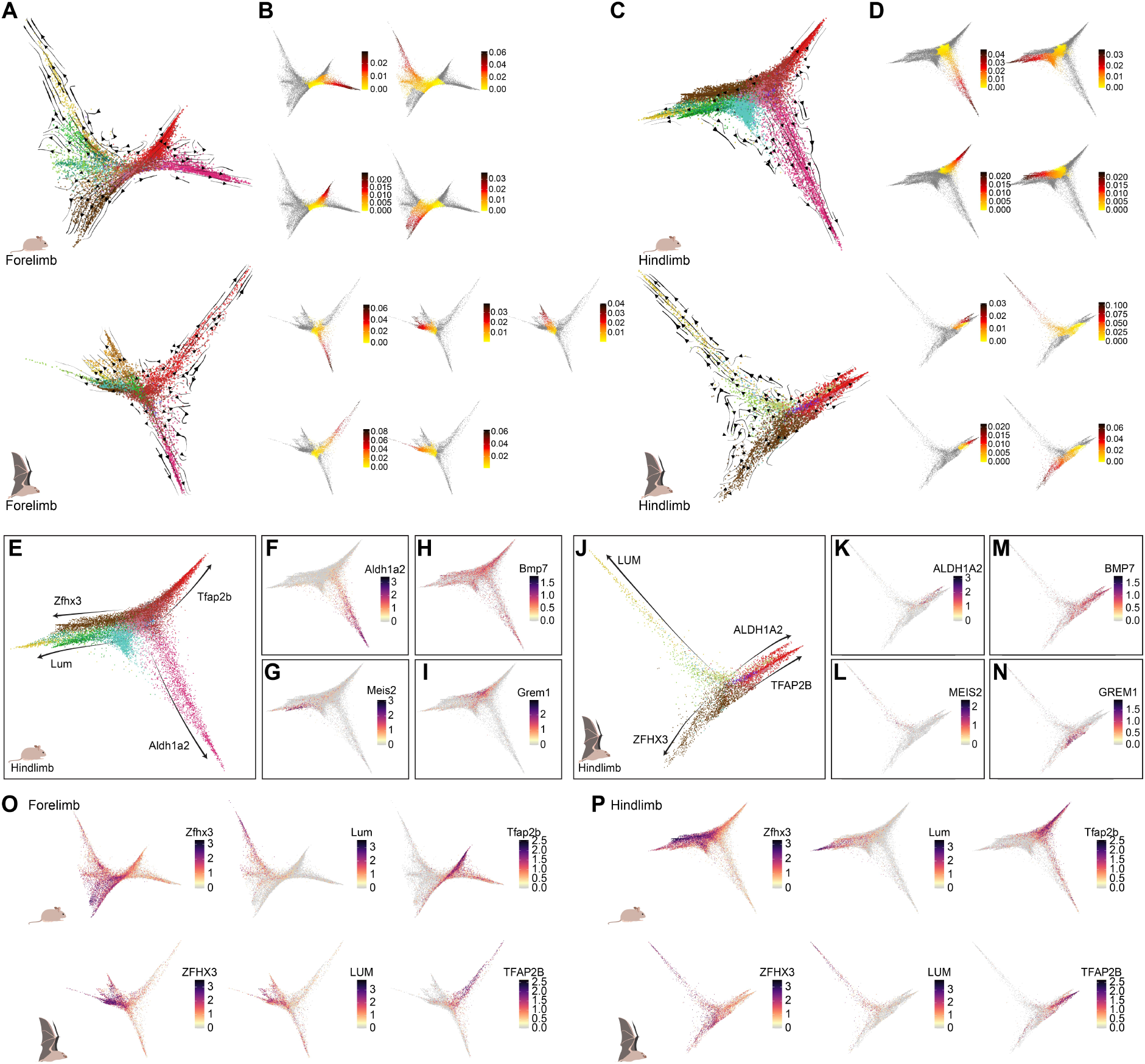
Differentiation trajectories of distal limb cells. **A** Combined CellRank kernels (0.2 * Pseudotime, 0.4 * Connectivity, 0.4 * Pseudotime) showing transition probabilities between non-chondrogenic Hoxd13+ cells of FLs of mouse (top) and bat (bottom). **B** Pseudotime values per trajectory as calculated using slingshot for FLs of mouse (top) and bat (bottom). **C** Combined CellRank kernels showing transition probabilities between non-chondrogenic *Hoxd13*+ cells of HLs of mouse (top) and bat (bottom). **D** Pseudotime values per trajectory as calculated using slingshot for HLs of mouse (top) and bat (bottom). **E** Differentiation trajectories of *Hoxd13*+, non-chondrogenic cells of the mouse hindlimb, derived from RNA velocity and pseudotime data indicated by arrows. Trajectories were annotated based on increasing expression of marker genes. **F-I** Shown is the expression of *Aldh1a2*, *Meis2*, *Bmp7* and *Grem1* in mouse HL trajectories. **J** Differentiation trajectories of *HOXD13*+, non-chondrogenic cells of the bat HL, derived from RNA velocity and pseudotime data indicated by arrows. Trajectories were annotated based on increasing expression of marker genes. **K-N** Shown is the expression of *ALDH1A2*, *MEIS2*, *BMP7* and *GREM1* in bat FL trajectories. **O** and **P** expression pattern of the genes used to annotate the differentiation trajectories in FLs (O) and HLs (P).

**Supplementary Figure 5. Related to Fig. 2.**
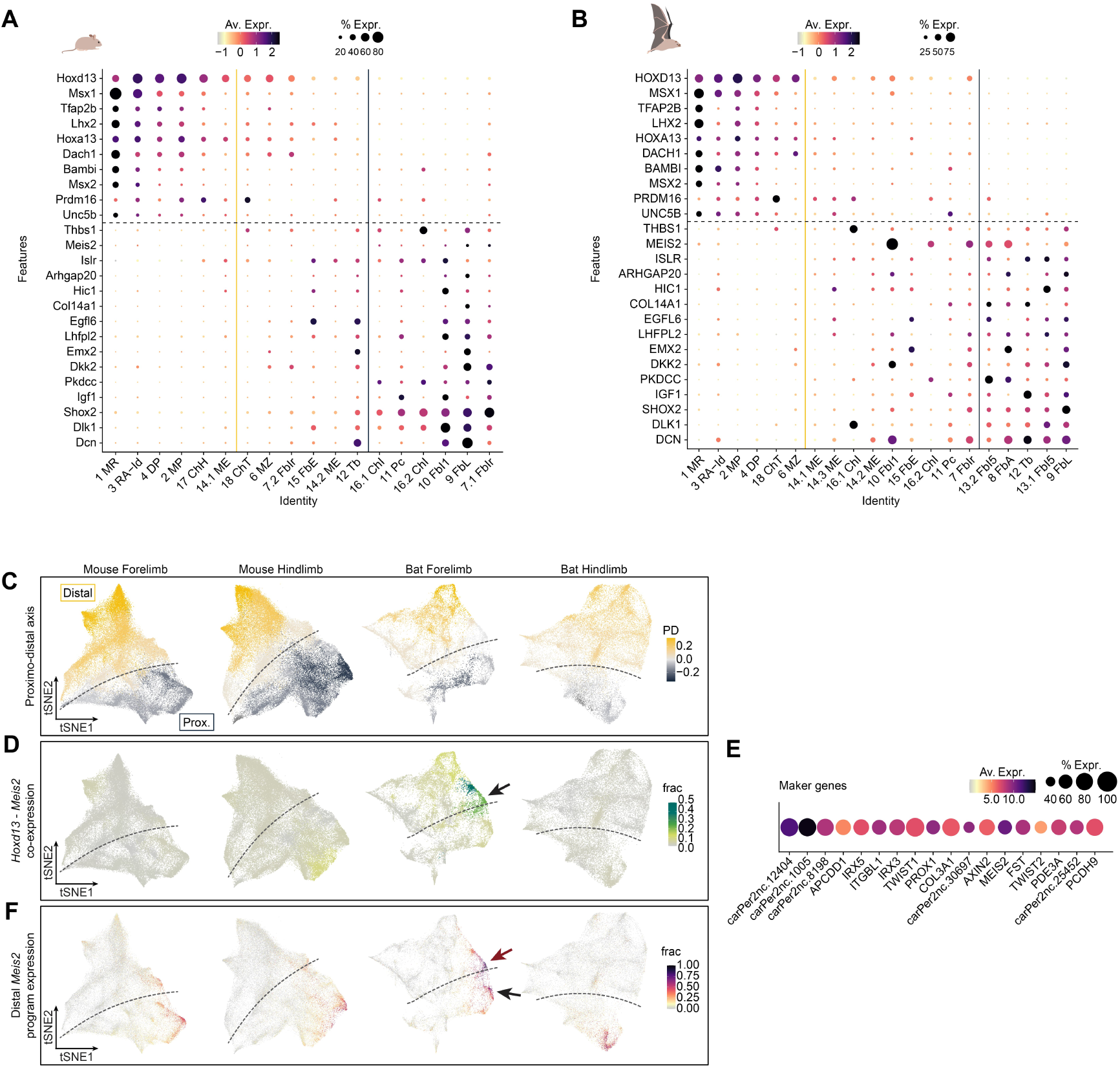
Proximo – Distal digital dissection. **A** Expression in mouse FLs of the genes found to be markers of FL distal mesenchymal cells, and the top 15 markers of FL proximal mesenchymal cells. Genes are ordered from top to bottom from most distal to most proximal. Clusters in the same order from left to right. **B** Expression in bat FLs of the genes found to be markers of FL distal mesenchymal cells, and the top 15 markers of FL proximal mesenchymal cells. **C** Assignment of a proximal (dark blue) or distal (yellow) identity to each cell of mouse and bat fore- and hindlimbs based on *Hoxd13* + *Msx1* and *Shox2* expression per cell. Shown are the differences in the proportion of proximally or distally assigned cells per cluster. **D** Co-expression of distal autopodial marker *Hoxd13* and chiropatagium marker *Meis2* in mouse and bat fore- and hindlimbs. Shown is the fraction of cells co-expressing both genes per cluster. Cluster 7 FbIr is highlighted with an arrow. **E** Marker genes of bat cluster 7 FbIr based on differential gene expression between cluster 7 and the rest of the LPM-derived bat FL cells. **F** Bat cluster 7 FbIr gene set expression in mouse and bat fore- and hindlimbs. Shown is the fraction of co-expression score of the whole marker gene set shown in E.

**Supplementary Figure 6. Related to Fig. 3A.**
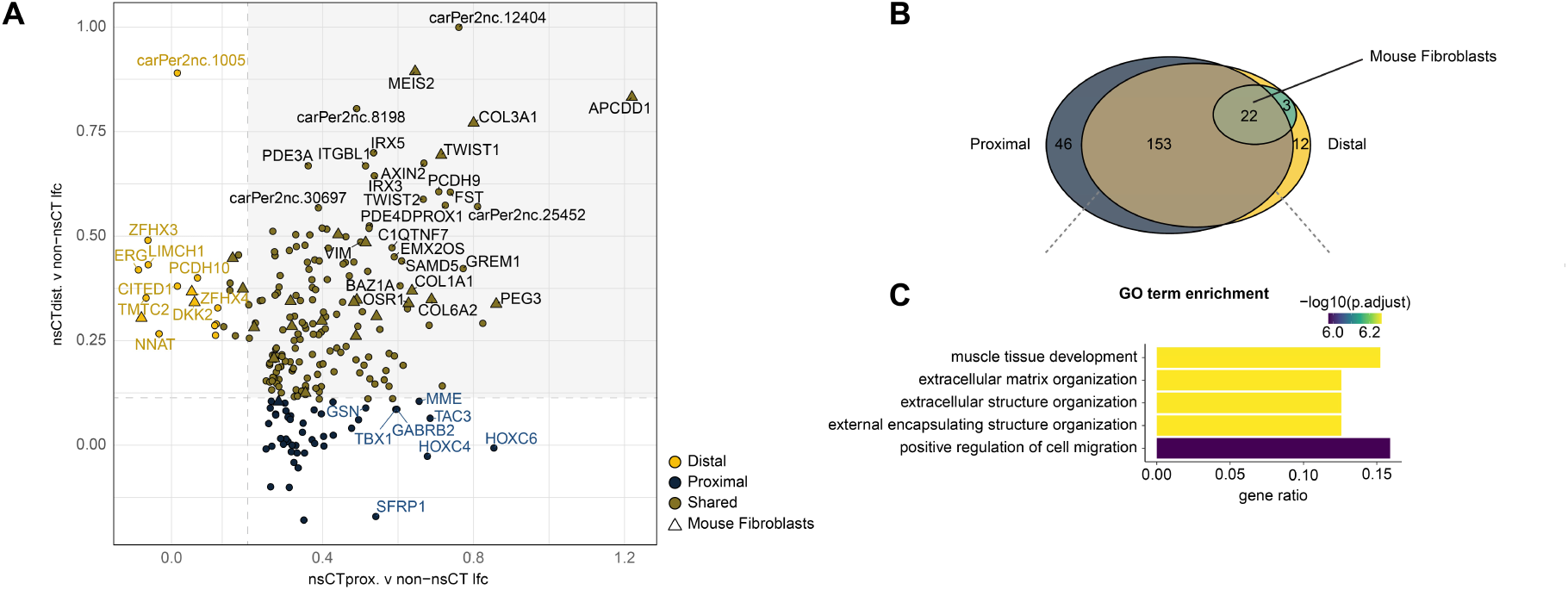
Second proximal gene program in the distal bat forelimb. **A** Correlation between the differential expression of genes from distal (cluster 7) and proximal (cluster 8) *MEIS2*- positive clusters in the bat forelimb identified in Extended Data Fig. 5. Shown is the logfold change of differential gene expression analysis of the respective cluster versus non-fibroblast LPM-derived cells. Genes shared with mouse fibroblasts are depicted as triangles. **B** Venn diagram showing the overlap (brown) between the genes enriched in the proximal (dark blue) and distal (yellow) cell subset as well as the fraction of genes shared with mouse fibroblasts (green). **C** GO term enrichment analysis of the 175 shared genes. Shown are the top 5 enriched GO terms.

**Supplementary Figure 7. Related to Fig. 3E.**
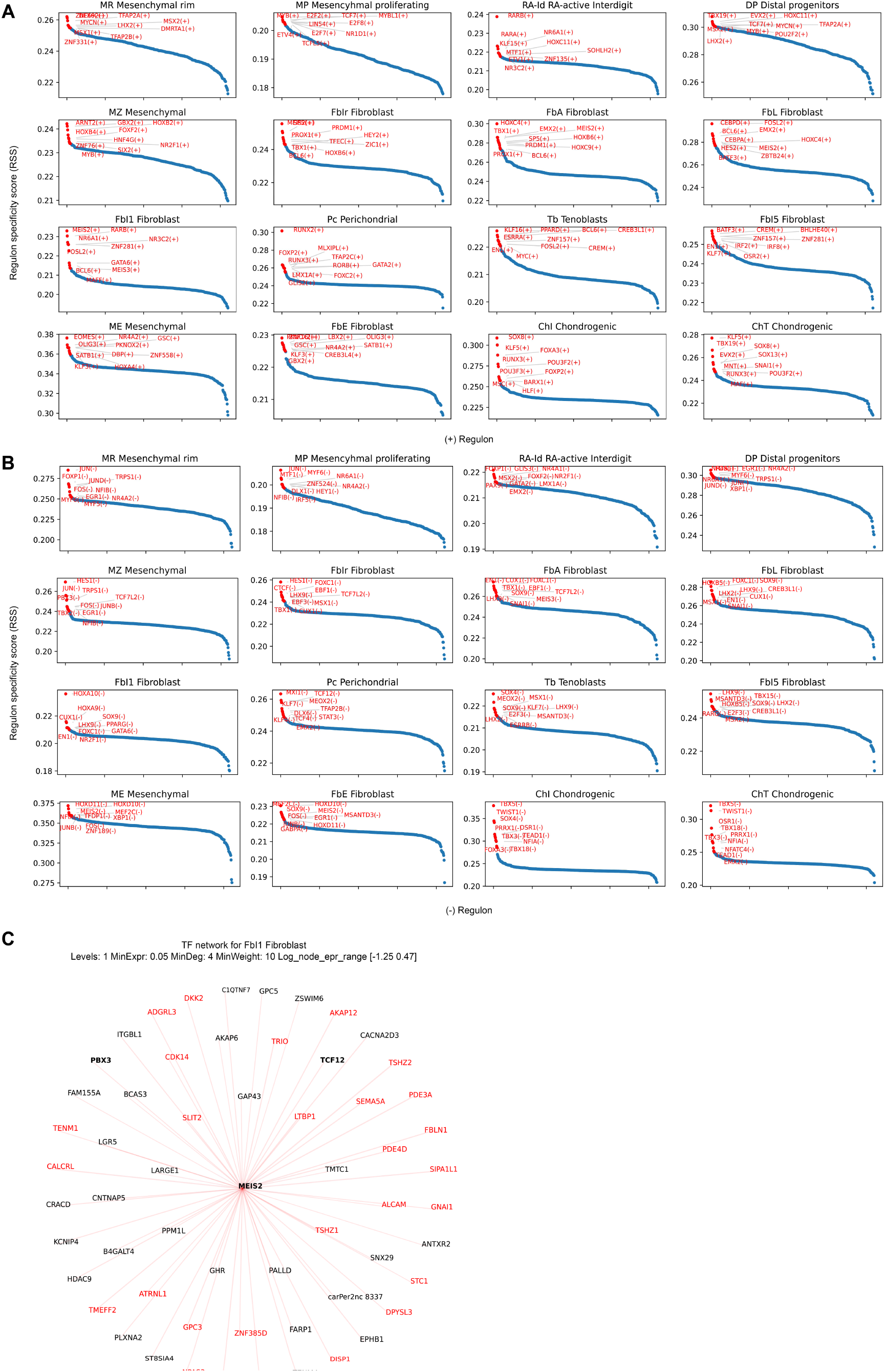
**A** and **B** Positive (A) and negative (B) regulons from the gene regulatory network analyses with SCENIC. **C** Transcription factor network showing downstream regulators for MEIS2 in bat forelimb cluster 10 (Fbl1).

**Supplementary Figure 8. Related to Fig. 3.**
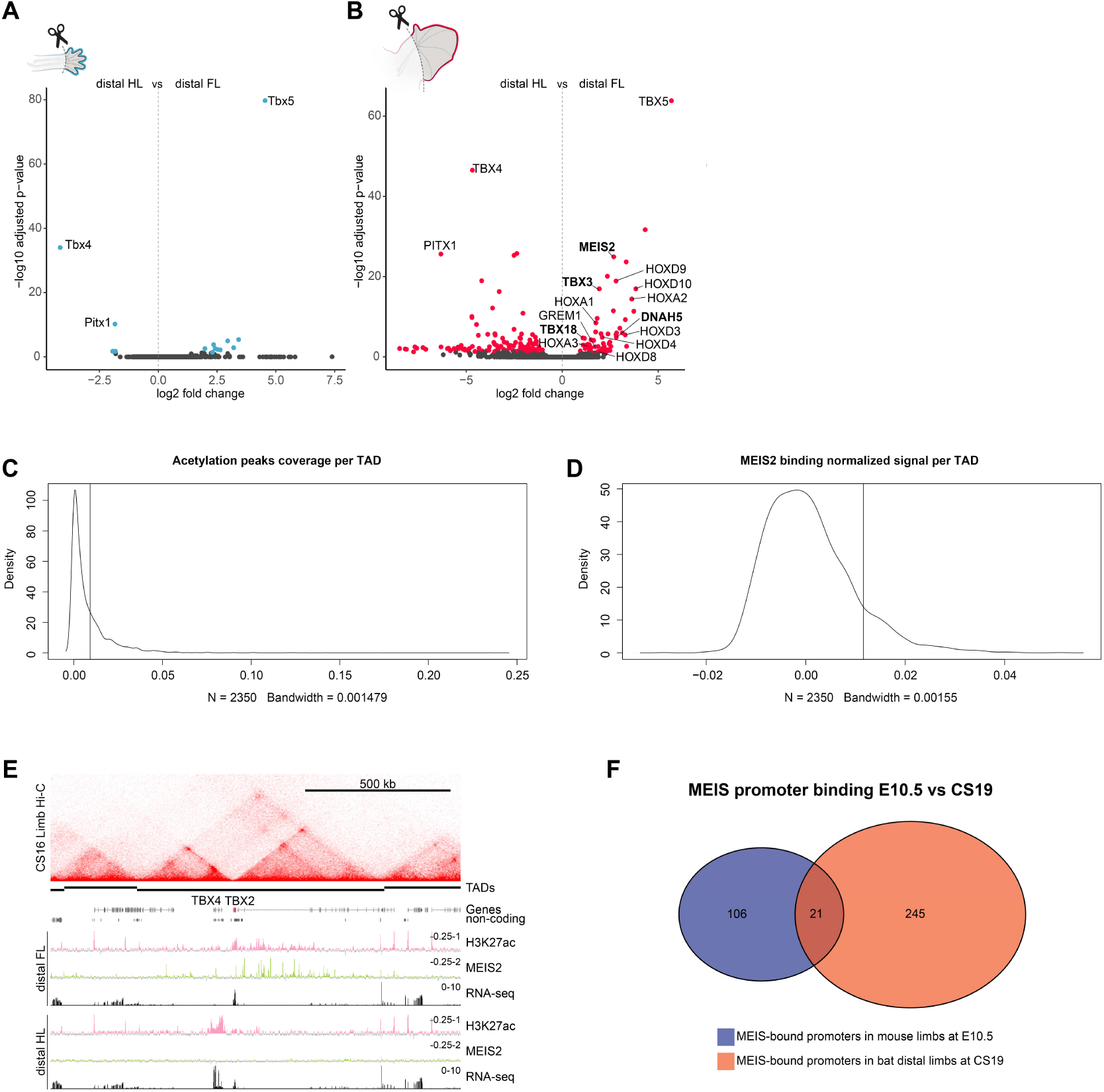
Epigenetic and transcriptomic bulk analyses. **A** and **B** Volcano plot of the differentially expressed genes from bulk RNA-seq between the dissected distal FL and distal HL in mice (A) and in bats (B). **C** Distribution of the TADs in the Bat genome, and the fraction of each of them covered by H3K27ac ChIP peaks specific to the distal FL. The vertical line shows the cutoff used to determine the enriched TADs. **D** Distribution of the TADs in the Bat genome, and the normalized (input subtracted) signal from MEIS binding found within each of them. The vertical line shows the cutoff used to determine the enriched TADs. **E** Bat *TBX4*/*TBX2* locus with Hi-C from CS16 FLs on top, TAD calling below. The Input subtracted H3K27ac ChIP-seq track is depicted in pink and input subtracted MEIS2 ChIP-seq track is shown in green. RNA-seq tracks are shown in black. ChIP-seq and RNA-seq were performed on distally dissected fore- and hindlimbs at CS18/19. Note the specific expression and acetylation of TBX4 in the HL and the specific MEIS2 binding sites throughout the *TBX2* domain unique to the distal FL. **F** Venn diagram showing the overlap (21) of MEIS-bound promoters in mouse 10.5 limbs and bat CS19 distal limbs.

**Supplementary Figure 9. Related to Fig. 4.**
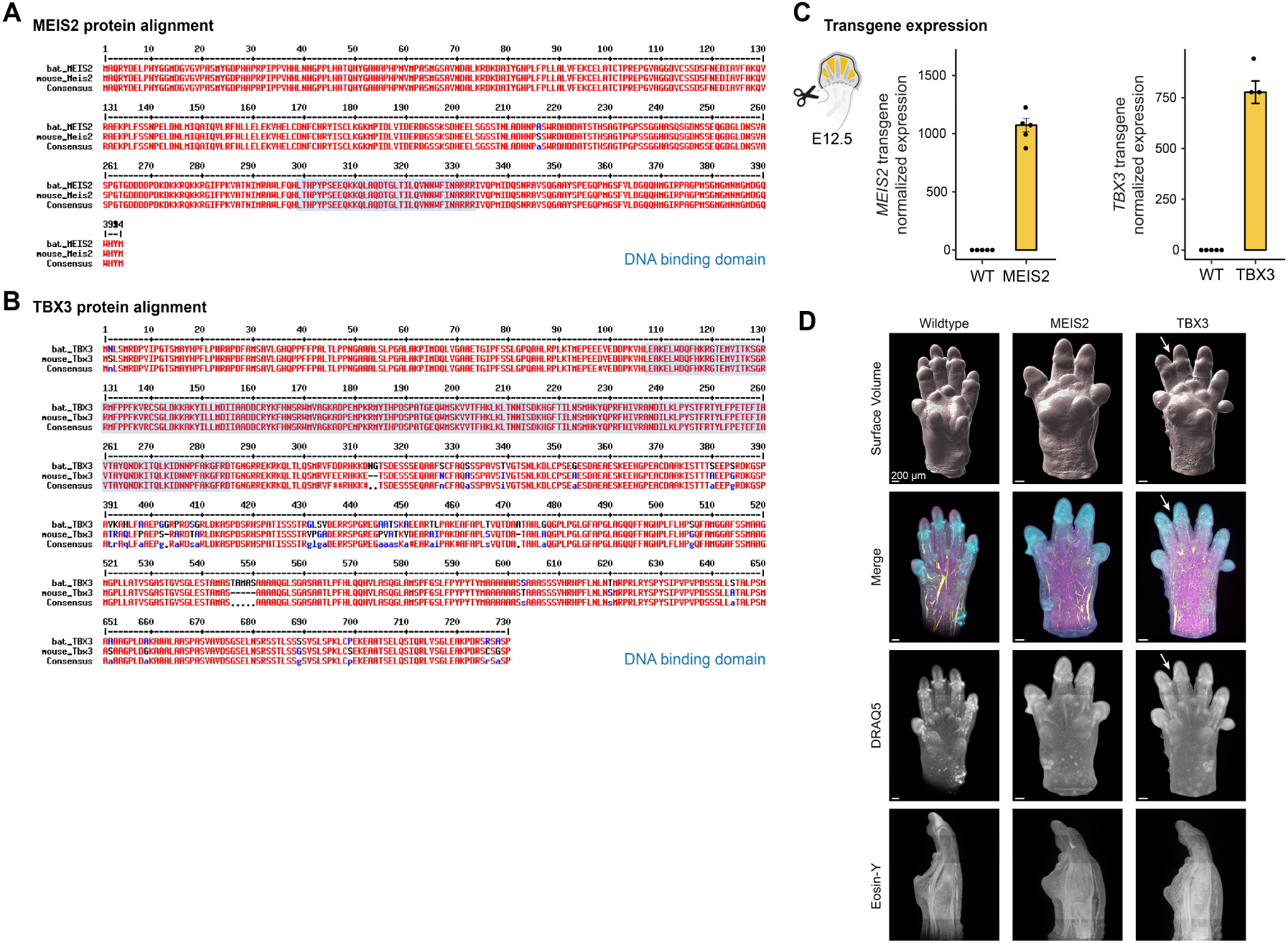
MEIS2 and TBX3 protein alignment, transgene expression and 3D imaging of mutant limbs. **A** and **B** Alignment of bat and mouse MEIS2 and TBX3 protein sequences. The DNA-binding domains are highlighted in blue. **C** RNA-seq of distally dissected wildtype and mutant limbs at E12.5 showing the normalized expression of *MEIS2* and *TBX3* transgenes. n=5. **D** 3D imaging of mouse wildtype and mutant whole limbs at E15.5. Shown is a surface representation, Eosin-Y staining and a merged imaged with an arrow highlighting syndactyly of digit II and III in the *TBX3* mutant.

